# Divergent spatial codes in retrosplenial cortex and hippocampus support multi-scale representation of complex environments

**DOI:** 10.1101/2024.08.22.609122

**Authors:** Célia Laurent, Nada El Mahmoudi, David M. Smith, Francesca Sargolini, Pierre-Yves Jacob

## Abstract

In everyday life, mammals navigate through environments composed of multiple interconnected spaces. Understanding how the brain encodes space in such complex settings is essential to uncover the mechanisms that support flexible navigation in the real world.

In this study, we investigated how the retrosplenial cortex (RSC) and the hippocampus (HPC) encode spatial information in multi-room environments by recording neuronal activity in rats exploring environments composed of two or four connected rooms. These environments varied both in their structural layout and in their sensory features, allowing us to disentangle the influence of environmental geometry from that of local sensory cues.

We found that two types of directional coding coexist within the RSC: classical head direction cells maintained a stable preferred firing direction across all rooms, while multidirectional cells expressed room-specific directional tuning that followed the geometric structure of the environment (e.g., 180° or 90° rotations). In addition, non-directional RSC neurons displayed room-specific spatial firing patterns that repeated across rooms following geometric transformations, similar to the activity observed in multidirectional cells. In contrast, hippocampal place cells either remapped between rooms or showed simple translational repetition, without preserving any geometric alignment.

Taken together, these findings reveal a functional dissociation between retrosplenial and hippocampal spatial codes. The RSC supports a structured, multi-scale representation of space, segmenting the environment into locally anchored reference frames embedded within a coherent global geometry. The HPC, by contrast, encodes room-specific representations independently for each compartment. This division may support flexible navigation in complex environments by integrating geometry-based spatial segmentation and unification in the RSC with context-specific encoding in the HPC, thereby enabling multi-level spatial coding across interconnected spaces.

## Introduction

Understanding how the brain encodes space in complex environments is essential to uncover the mechanisms that support flexible navigation under naturalistic conditions. However, while most models of spatial representation have been developed in simple, open environments, real-world spaces typically consist of multiple interconnected rooms. The structural divisions of these environments impose discrete transitions that disrupt the continuity of sensory and spatial experience. This raises the possibility that the brain actively segments space into distinct yet connected representations in order to maintain coherent spatial orientation.

The retrosplenial cortex (RSC) is a strong candidate for supporting such spatial segmentation. Functionally, it is involved in navigation, memory, and contextual processing (Takahashi *et al*., 1997; Sherrill *et al*., 2013; Mitchell *et al*., 2018; Miller *et al*., 2019; Vedder *et al*., 2016). Anatomically, the RSC is well positioned to integrate spatial and sensory information, through connections to the medial entorhinal cortex and the hippocampus (HPC), two critical structures for spatial cognition (Cenquizca & Swanson, 2007; Czajkowski *et al*., 2013; Jones & Witter, 2007; Naber *et al*., 2001; Sugar & Witter, 2016; Van Groen & Wyss, 1990, 2003). The RSC also receives direct input from primary visual areas (Vélez-Fort *et al*., 2018; Yoder *et al*., 2011; Wyss & Van Groen, 1992, 2003), which is consistent with a role in incorporating visual landmarks into spatial representations (Sit and Goard, 2023). Taken together, these studies suggests that the RSC may serve as an integrative hub linking local sensory cues, directional orientation, and contextual memory.

Recent studies have identified multidirectional cells (MDCs) in the RSC, a neuronal population that emerges specifically in multicompartment environments (Jacob *et al*., 2017; Zhang *et al*., 2022). Unlike classical head direction cells (HDCs), which maintain a single preferred firing direction (PFD) across different rooms, MDCs exhibit a distinct PFD in each room. This produces bidirectional tuning in two-room environments and tetradirectional tuning in four-room layouts. These findings suggest that the RSC could support both global and local spatial reference frames, encoded by HDCs and MDCs respectively.

However, the functional significance of MDCs remains unclear. Previous studies have proposed that MDCs tuning reflects anchoring to local sensory cues or to global symmetry, such as the rotational repetition of identical rooms (Jacob *et al*., 2017; Zhang *et al*., 2022). However, as these environments used visually indistinguishable rooms, it is still difficult to determine whether MDCs reflect true structural segmentation or arise from perceptual ambiguity. Furthermore, these studies did not investigate whether directional tuning is organized according to the geometry of the environment, which is defined here as the spatial layout and angular relationships between rooms. Finally, it is unclear whether such geometry-aligned activity is exclusive to directionally tuned neurons or implies a more general coding principle across the RSC.

Recent work has shown that the RSC encodes spatial context through distributed population activity (Sun *et al*., 2021; Miller *et al*., 2021; Trask & Helmstetter, 2022; Subramanian *et al*., 2024), suggesting that spatial segmentation may extend beyond directional tuning. However, these studies were conducted in simple, open arenas and did not address whether contextual or structural segmentation generalizes to geometrically structured multi-room environments.

To address these questions, we recorded neuronal activity in rats navigating in environments consisting of two or four interconnected rooms, which were either visually and tactilely identical or distinct. This design allowed us to distinguish between the influence of sensory cues and that of environmental geometry on spatial tuning. We analyzed the activity of HDCs, MDCs, and non-directional RSC neurons to determine whether spatial segmentation generalizes across neuronal subtypes, and whether it reflects structural layout rather than local perceptual inputs. In parallel, we recorded the activity of hippocampal place cells under identical conditions to test whether the RSC and the HPC rely on shared or divergent coding principles.

Our results demonstrate that the RSC encodes a structured representation of space that is aligned with both room segmentation and global environmental geometry. HDCs provide a stable, environment-wide directional signal, whereas MDCs and non-directional neurons exhibit room-specific tuning patterns that undergo predictable geometric transformations across rooms: reversing by 180° in two-room environments and rotating by 90° in four-room layouts. These geometry-aligned codes remained consistent throughout sensory changes and stable over time, suggesting that spatial structure, rather than local cues, organizes RSC representations. In contrast, hippocampal place cells remapped between rooms without preserving any consistent spatial alignment, suggesting a distinct, context-dependent coding strategy. Together, these findings demonstrate that the RSC supports a flexible, geometry-aligned segmentation of space that generalizes across cell types, providing a neural scaffold for encoding complex environments as discrete spatial units.

## Results

### 1. Multidirectional cells (MDCs) encode room-specific directional codes

#### 1.1 Bidirectional tuning is independent of perceptual features

To investigate the properties of multidirectional cells (MDCs) in complex environments, we implanted recording electrodes in the dysgranular retrosplenial cortex (**Supplementary Figure 2**). In two-room settings, MDCs typically exhibit two preferred directions and are referred to as bidirectional cells (BDCs) (Jacob *et al*.,2017). First, we identified 30 BDCs recorded in 9 rats, navigating in a two-room environment with visually identical rooms (**S1-2id**; **Figure 1A-C, top**). The distribution of preferred firing directions (PFDs) was uniform (**Supplementary Figure 3**; Rayleigh test, Z = 0.45, p = 0.64), indicating that BDCs spanned the full 360° directional space uniformly.

**Figure 1.**
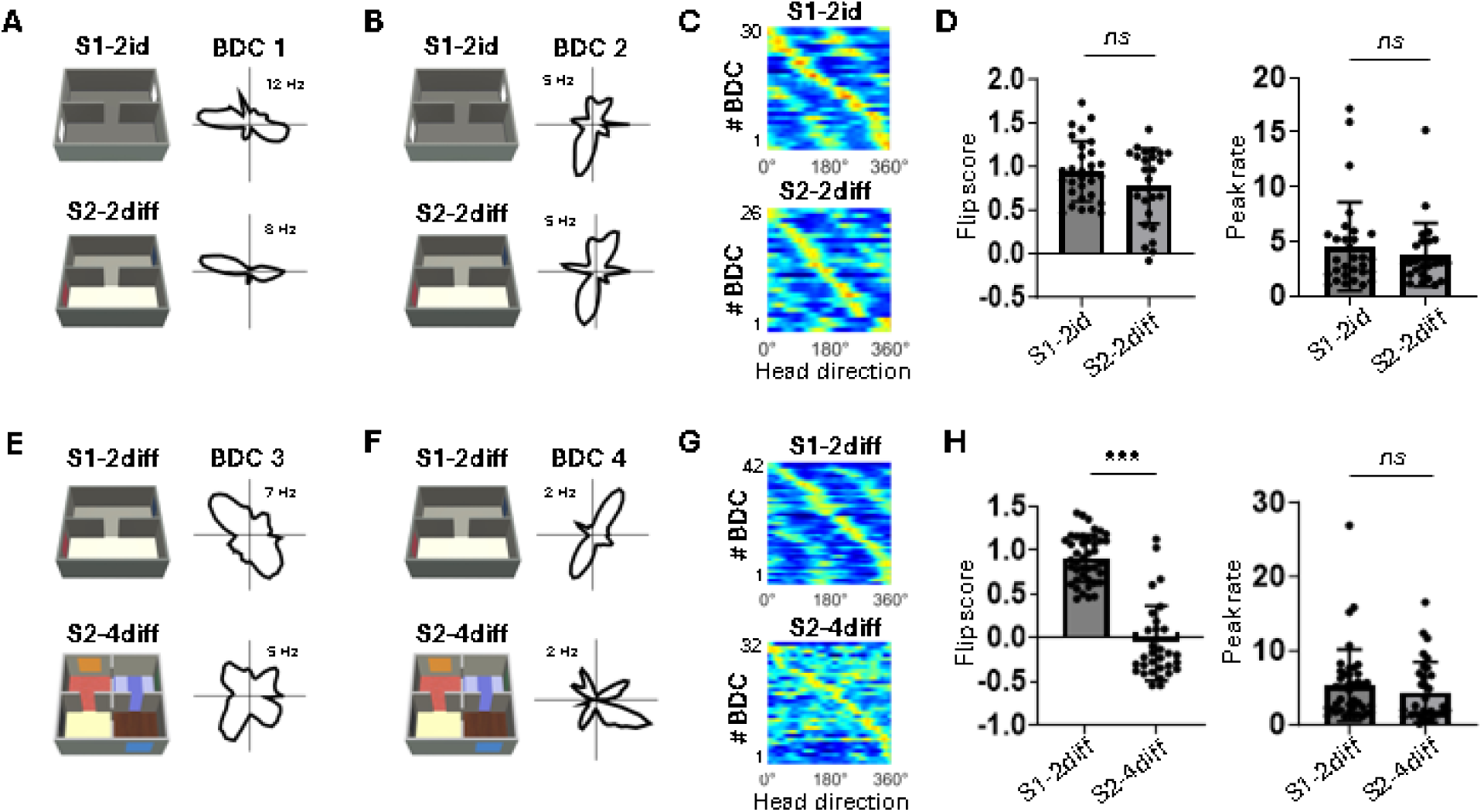
Multidirectional cells (MDCs) exhibit stable tuning across perceptual changes and adapt to environmental complexity. **A-B.** *Left*: Animals successively explored two environments composed of two connected rooms that were either visually and tactilely identical (S1–2id: same visual cue and floor) or distinct (S2–2diff: different visual cues and floors). *Right:* Polar plots of two example BDCs (BDC1 and BDC2) recorded in both conditions. Peak firing rates are indicated for each cell. Note the emergence of secondary peaks along the 180° axis, regardless of the perceptual similarity between rooms. **C.** Directional activity of 30 BDCs recorded in S1–2id (top); among them, 26 were also recorded in S2–2diff (bottom). Activity is color-coded from blue (no activity) to orange (peak firing rate), and cells are sorted by their PFD. **D.** Comparison of BDC properties between S1–2id and S2–2diff. *Left:* Flip scores, quantifying bidirectionality is performed by computing the difference at 180° *vs*. 90° of the tuning curve autocorrelation. *Right:* Peak firing rates. No significant differences were observed between environments. Bars represent mean ± SD; individual data points are shown as black dots. Statistical comparisons were performed using unpaired t-tests. **E-F.** *Left:* Animals successively explored two environments composed of either 2 (S1–2diff) or 4 (S2–4diff) visually and tactilely distinct connected rooms *Right:* Polar plots of two example BDCs (BDC3 and BDC4) recorded in both conditions. Note the emergence of multiple peaks in the four-room condition. **G.** Directional activity of 42 BDCs recorded in S1– 2diff (top); 32 of these were also recorded in S2–4diff (bottom). Color scale and sorting as in panel C. **H.** Comparison of BDC properties between S1–2diff and S2–4diff. *Left:* Flip scores, computed as in D, significantly decreased in the four-room condition, indicating a loss of bidirectionality. *Right:* Peak firing rates did not change significantly between environments. Bars represent mean ± SD individual data points are shown as black dots. Statistical comparisons were performed using unpaired t-tests. ***p < 0.0001.

To test whether bidirectional tuning depends on perceptual similarity between rooms, we re-exposed the same animals to a structurally identical environment in which the two rooms were now distinguished by salient visual and tactile cues (**S2-2diff**; **Figure 1A-C, bottom**). Of the 26 BDCs recorded in both settings, the majority (77%) maintained two opposing peaks of activity. Quantitative measures showed no significant changes in flip score (**Figure 1D-left:** unpaired t-test, t(54) = 1.62, p = 0.11), double-angle vector length (**Supplementary figure 4**: unpaired t-test, t(54) = 1.53, p = 0.13), or peak firing rate (**Figure 1D-right**: unpaired t-test, t(54) = 0.81, p = 0.42). Directional tuning also remained uniformly distributed (**Supplementary figure 3**: Rayleigh test, Z = 1.27, p = 0.28).

These findings indicate that bidirectional tuning in BDCs is preserved despite salient perceptual differences between rooms. This suggests that the directional coding in these cells may be primarily driven by spatial configuration rather than sensory cues.

#### 1.2 BDC tuning becomes multidirectional with increasing environmental complexity

To test whether tuning adapts to increasing spatial complexity, we observed 32 BDCs recorded in 2 additional rats as they transitioned from a two-room environment (**S1-2diff**) to a four-room configuration (**S2-4diff**), ensuring that all rooms were perceptually distinct (**Figure 1E-G**). Rayleigh tests on the PFDs of the BDCs revealed no significant clustering in either environment (**Supplementary Figure 3**: Rayleigh tests for S1-2diff and for S2-4diff: Z = 0.87, p = 0.42 and Z = 1.41, p = 0.25, respectively), indicating that BDCs covered the full 360° directional space.

In the S2-4diff condition, BDCs exhibited additional tuning peaks, indicating a shift from bidirectional to multidirectional activity. This transformation was reflected in a significant drop in the flip score (**Figure 1H-left**: unpaired t-test, t(72) = 11.97, p < 0.001), while the peak firing rate remained stable across conditions (**Figure 1H-right**: unpaired t-test, t(72) = 0.90, p = 0.37).

These results show that BDCs develop multidirectional tuning in environments comprising multiple rooms, suggesting that their directional code adapts to spatial complexity. This pattern indicates a directional code that reflects the structural segmentation of space.

#### 1.3 Inter-room tuning shifts reflect environmental geometry

To further examine how tuning patterns relate to environmental structure, we analyzed PFD stability within and across rooms. In both two-room environments (S1-2id and S2-2diff, **Figure 2A-C**), BDCs showed unipolar tuning within each room (**Supplementary Figure 6A**: S1-2id, paired t-test, t(29) = 0.27, p = 0.79; S2-2diff, paired t-test, t(25) = 0.28, p = 0.78).

**Figure 2.**
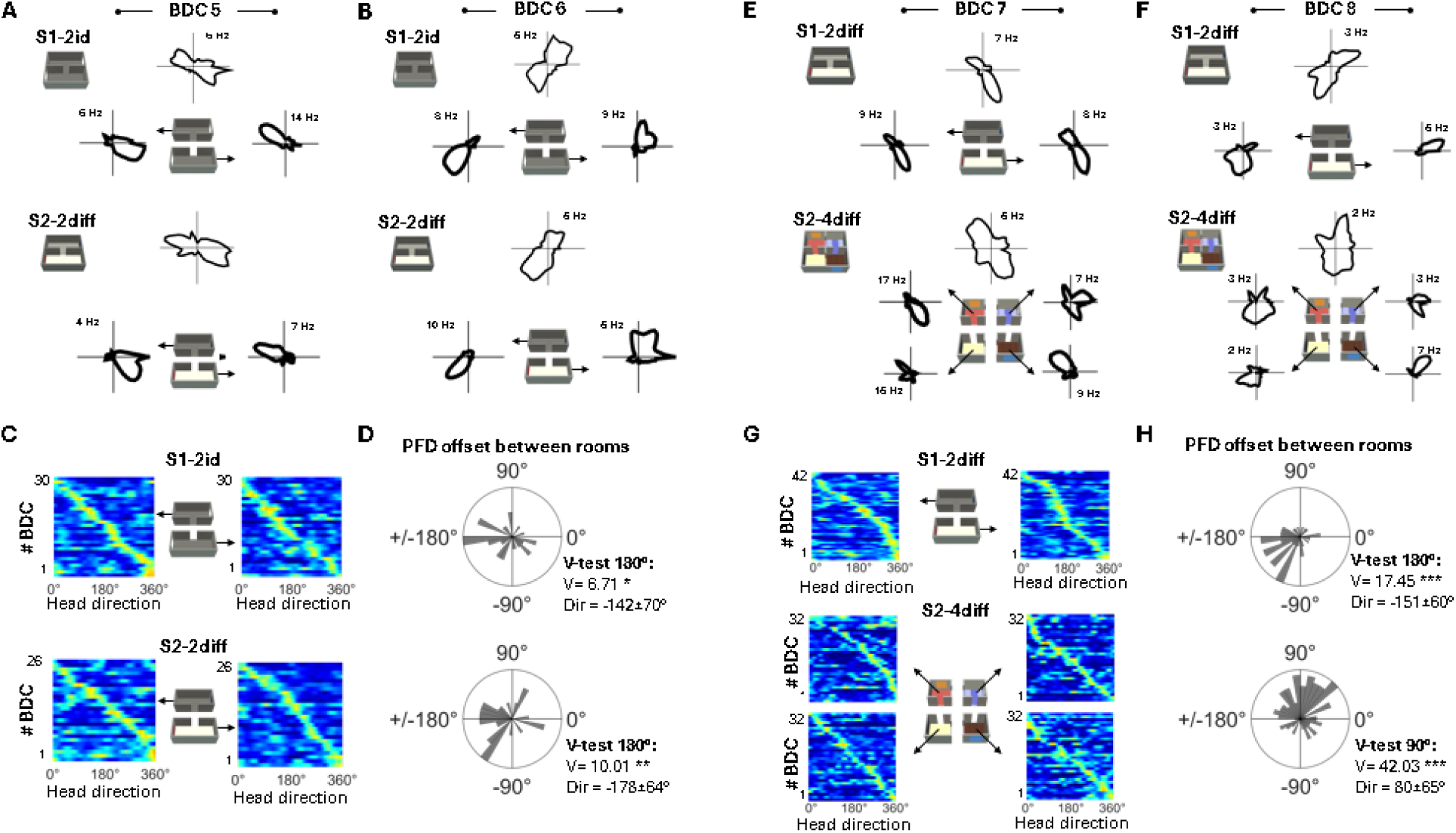
Inter-room tuning shifts of multidirectional cells (MDCs) reflect environmental geometry. **A-B.** Animals successively explored two environments composed of two connected rooms that were either visually and tactilely identical (S1-2id) or distinct (S2-2diff). Polar plots of two example BDCs (A: BDC5; B: BDC6) are shown, plotted separately for each room. Both cells exhibited unipolar activity within individual rooms, regardless of perceptual similarity. **C.** Directional activity for each room of 30 BDCs recorded in S1-2id (top); 26 of these were also recorded in S2-2diff (bottom). All cells displayed unipolar tuning in each room. **D.** Polar histograms showing angular differences in PFD between rooms in S1-2id (top) and S2-2diff (bottom). Distributions cluster significantly around 180°, indicating opposite tuning directions across rooms. V: V-test; Dir: mean PFD offset between rooms ± SD. ***p < 0.0001. **E-F.** Animals successively explored two environments composed of either 2 (S1–2diff) or 4 (S2–4diff) visually and tactilely distinct connected rooms. Polar plots of two example BDCs (E: BDC7; F: BDC8), shown separately for each room. Both cells exhibited unipolar activity within individual rooms, even when the global pattern appeared multidirectional. **G.** Directional activity for each room of 42 BDCs recorded in S1-2diff (top); 32 of these were also recorded in S2-4diff (bottom). All cells showed unipolar tuning within each room. **H.** Polar histograms showing angular differences in PFD between rooms for S1-2diff (top) and S2-4diff (bottom). Distributions cluster significantly around 180° in the two-room condition and around 90° in the four-room condition, consistent with the spatial layout. Peak frequencies are indicated on each polar plot. V: V-test; Dir: mean PFD offset between rooms ± SD. ***p < 0.0001.

Next, we examined whether inter-room shifts in directional tuning followed consistent spatial relationships by calculating angular offsets between rooms (**Figure 2D**). In both conditions, PFD differences clustered significantly around 180° (S1-2id, V-test for 180°: V = 6.71, p = 0.042; S2-2diff, V-test for 180°: V = 10.01, p = 0.003), indicating that BDCs reversed their tuning across rooms.

To determine whether this pattern generalized to more complex environments, we analyzed BDC activity within rooms of the S1-2diff and S2-4diff conditions. BDCs continued to exhibit stable unipolar tuning within each room (**Supplementary Figure 6B**: S1-2diff, paired t-test: t(41) = 0.10, p = 0.92; S2-4diff, one-way ANOVA: F(2.492, 77.25) = 0.73, p = 0.51). However, inter-room angular offsets varied systematically with environmental geometry: in the S1-2diff condition, PFDs differed by ∼180° again (**Figure 2H**: V-test for 180°: V = 17.45, p < 0.0001), while in the S2-4diff condition, offsets clustered significantly around 90° (V-test for 90°: V = 42.03, p < 0.0001), which is consistent with a tetradirectional activity.

These results indicate that BDCs maintain consistent unidirectional tuning within rooms and exhibit systematic shifts between rooms aligned with environmental geometry. This geometry-dependent modulation supports the idea of a directional code specific to individual rooms of MDCs.

### 2. Head direction cells (HDCs) encode a stable global reference frame

#### 2.1. HDCs tuning remains stable across perceptual changes

In parallel with the recordings of bidirectional cells, we identified 77 classical head direction cells (HDCs) in the two-room environment composed of two visually identical connected rooms (S1-2id; **Figure 3A-C**). The distribution of preferred firing directions (PFDs) was uniform (**Supplementary Figure 3**; Rayleigh test, Z = 0.62, p = 0.54), confirming that the HDCs covered the full 360° directional space uniformly (**Supplementary Figure 3**).

**Figure 3.**
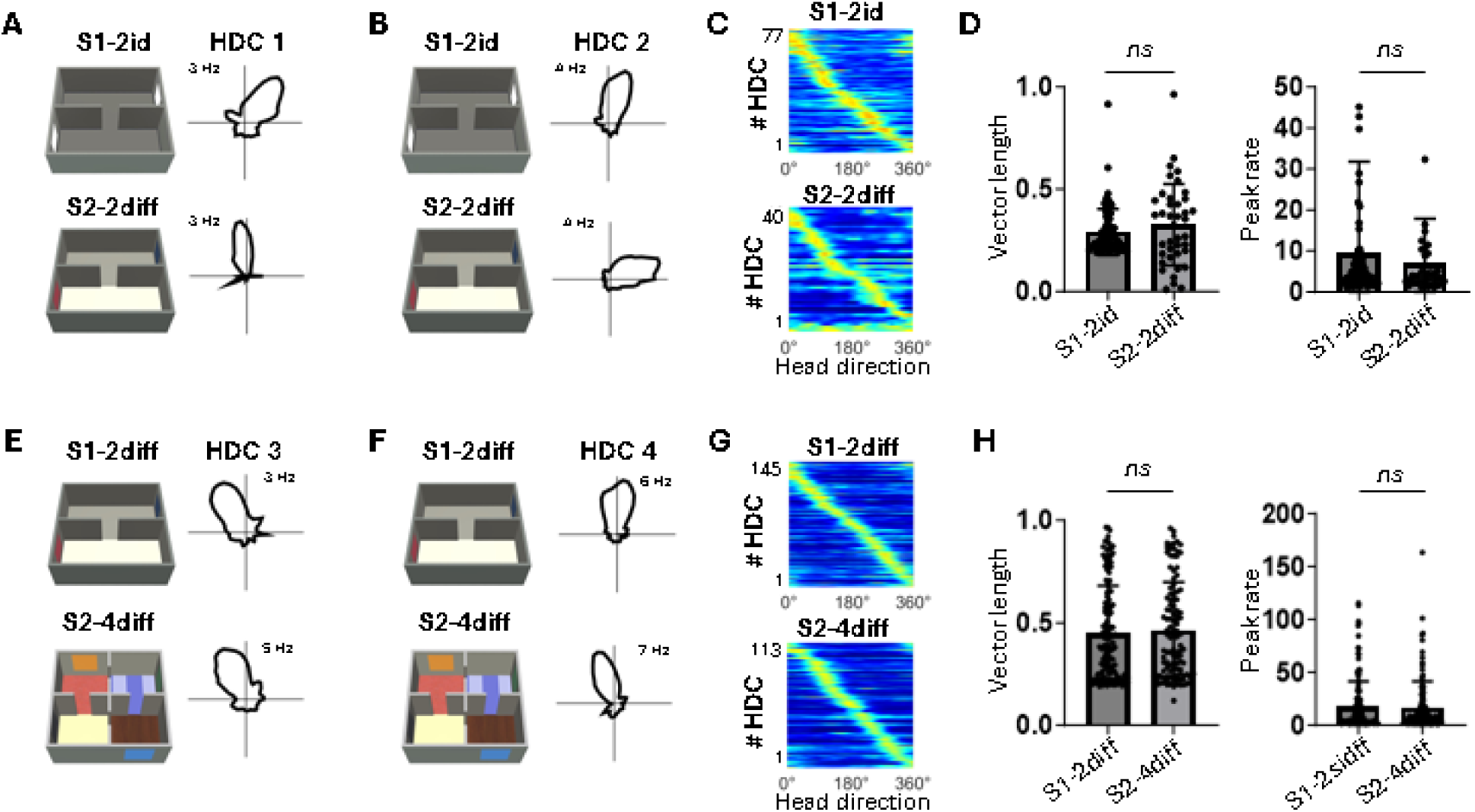
Head direction cells maintain stable directional tuning across perceptual and structural changes. **A-B**. *Left*: Animals successively explored two environments composed of either visually identical (S1–2id) or visually and tactilely distinct (S2–2diff) connected rooms. *Right*: Polar plots of two example HDCs (HDC1 and HDC2) recorded in both environments. Peak firing rates are indicated for each cell. **C**. Directional activity of the 77 HDCs recorded in S1–2id (top); among them, 40 were also recorded in S2–2diff (bottom). Activity is color-coded from blue (no activity) to orange (peak firing rate), and cells are sorted by preferred firing direction (PFD).**D**. Comparison of HDC properties between S1–2id and S2–2diff. *Left*: Mean vector length. *Right*: Peak firing rate. No significant differences were observed between environments. Bars represent mean ± SD; individual data points are shown as black dots. Statistical comparisons were performed using unpaired t-tests. **E-F**. *Left*: Animals successively explored environments composed of either two (S1–2diff) or four (S2–4diff) visually and tactilely distinct connected rooms. *Right*: Polar plots of two example HDCs (HDC3 and HDC4) recorded in both conditions. **G**. Directional activity of 145 HDCs recorded in S1-2id (top); among them, 113 HDCs were also recorded in S2-4diff (bottom). Activity is color-coded as in panel C, and cells are sorted according to their PFD. **H**. Comparison of HDC properties between S1–2diff and S2–4diff. *Left*: Mean vector length. *Right*: Peak firing rate. No significant differences were observed between environments. Bars represent mean ± SD; individual data points are shown as black dots. Statistical comparisons were performed using unpaired t-tests. ***p < 0.0001

To test the stability of their tuning in the face of perceptual changes, we followed 40 of these HDCs in a second environment (S2-2diff), which had the same layout but featured different visual and tactile cues in each room. Despite these differences, HDCs maintained their unidirectional tuning properties. Neither the Rayleigh vector length nor the peak firing rate changed significantly across environments (**Figure 3D**: unpaired t-tests for vector length, t(115) = 1.23, p = 0.22; unpaired t-tests for peak firing rate, t(115) = 0.64, p = 0.52). Directional tuning remained uniformly distributed in S2-2diff as well (Rayleigh test, Z = 0.79, p = 0.46), which further support the stability of the HDCs directional signal (**Supplementary Figure 3**).

These findings suggest that HDCs encode a stable global directional signal that generalizes across perceptually distinct but, structurally similar environments, indicating that their tuning is unaffected by local sensory identity.

Notably, some BDCs and HDCs were recorded simultaneously in both environments (**Supplementary Figure 5**), demonstrating that room-specific and environment-wide directional codes coexist within the retrosplenial cortex.

#### 2.2. HDCs maintain coherence in complex environments

To assess whether HDC tuning persists in more spatially complex settings, we recorded 145 HDCs in the two-room environment composed of visually and tactilely distinct rooms (S1-2diff), and then tracked 113 of these HDCs in the four-room configuration composed of equally distinct rooms (S2-4diff; **Figure 3E-G**).

Across these conditions, HDCs preserved their directional properties; there was no significant difference in either Rayleigh vector length or peak firing rate between the environments (**Figure 3H**: unpaired t-test for vector length, t(256) = 0.62, p = 0.53; unpaired t-test for peak firing rate t(256) = 0.22, p = 0.83). The distribution of PFDs remained uniform in S2-4diff (Rayleigh test, Z = 0.37, p = 0.69), though a modest clustering was observed in S1-2diff (Z = 3.21, p = 0.04; **Supplementary Figure 3**).

Once again, several HDCs were recorded simultaneously with BDCs in both conditions (**Supplementary Figure 5**), which provides further evidence for the parallel encoding of multiple directional codes within the RSC.

These results show that HDCs maintain a coherent unidirectional signal across segmented, perceptually diverse environments. Unlike MDCs, their tuning remains stable across rooms, consistent with a global directional reference frame.

#### 2.3. HDCs tuning remains consistent across room boundaries

Next, we examined whether HDCs preserve consistent tuning between rooms within the same environment. Across all four configurations (S1-2id, S2-2diff, S1-2diff, and S2-4diff), HDCs maintained highly similar preferred firing directions (PFDs) across rooms. In both S1-2id and S2-2diff, vector lengths remained comparable between rooms (**Supplementary Figure 6C**; S1-2id, paired t-test: t(76) = 0.41, p = 0.68; S2-2diff, paired t-test, t(39) = 0.52, p = 0.60). Angular offsets in PFDs were significantly clustered around 0° (**Figure 4D**; S1-2id, V-test for 0°: V = 48.05, p < 0.0001; S2-2diff, V-test for 0°: V = 20.79, p < 0.0001).

**Figure 4.**
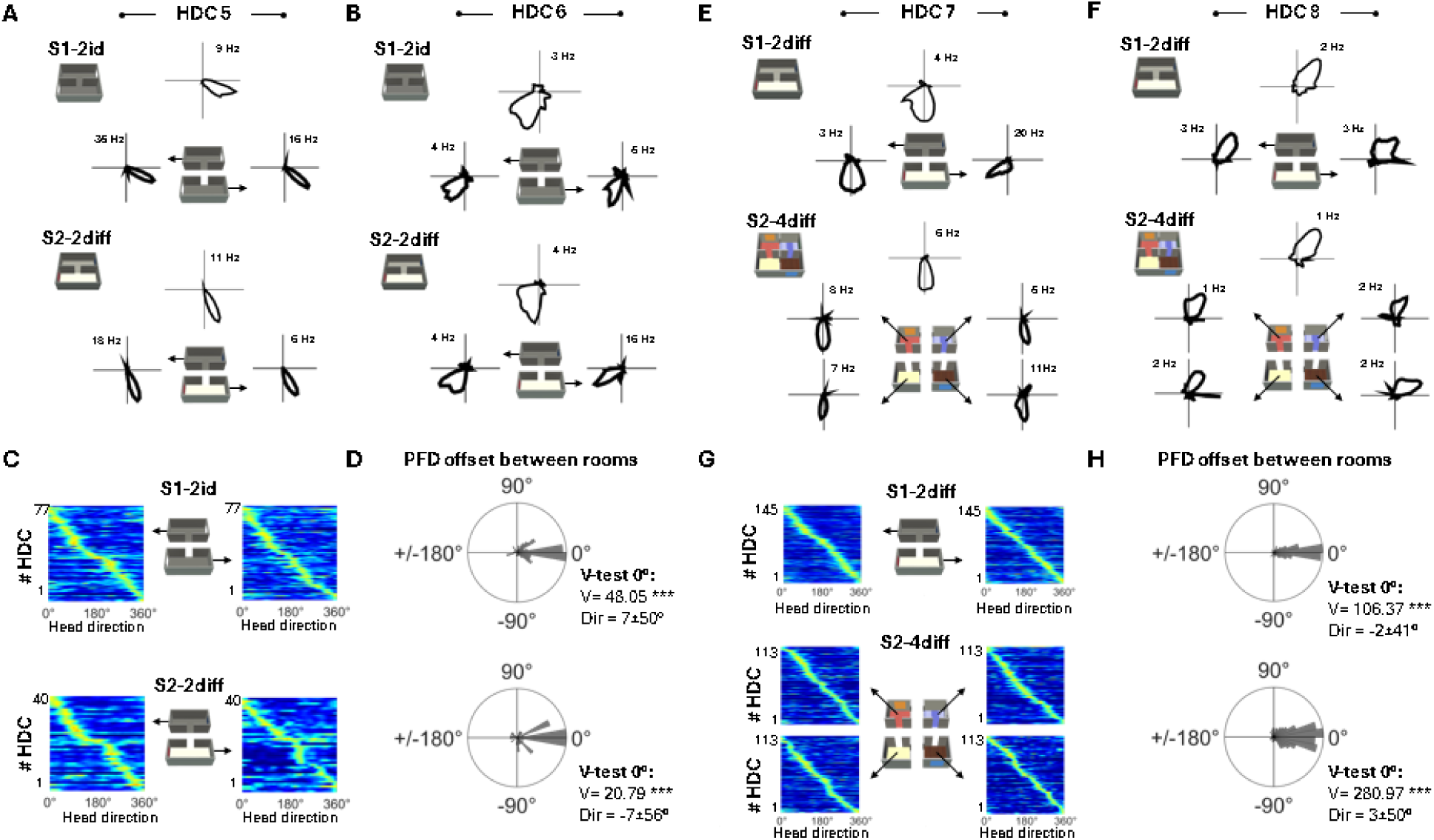
Head direction cell tuning remains consistent across room boundaries. **A-B**. Animals successively explored two environments composed of two connected rooms that were either visually and tactilely identical (S1-2id) or distinct (S2-2diff). Polar plots of two example HDCs (A:HDC5; B: HDC6) are shown separately for each room. Both cells exhibited the same unipolar activity in each room, regardless of perceptual similarity. **C**. Directional activity in each room of 77 HDCs recorded in S1-2id (top); 40 of these were also recorded in S2-2diff (bottom). All cells maintained unipolar tuning in each room. Activity is color-coded from blue (no firing) to orange (peak firing rate), and cells are sorted by PFD. **D**. Polar histograms showing angular differences in PFD between rooms in S1–2id (top) and S2– 2diff (bottom). In both environments, PFDs cluster significantly around 0°, indicating stable directional tuning across rooms. V: V-test; Dir: mean PFD offset between rooms ± SD. ***p < 0.0001. **E-F**. Animals successively explored environments composed of either two (S1– 2diff) or four (S2–4diff) visually and tactilely distinct connected rooms. Polar plots of two example HDCs (E: HDC7; F: HDC8) are shown separately for each room. Both cells maintained consistent unipolar activity across all rooms, regardless of the number of connected rooms. **G**. Directional activity in each room of 145 HDCs recorded in S1-2diff (top); 113 of these were also recorded in S2-4diff (bottom). All cells exhibited consistent unipolar tuning in each room. **H**. Polar histograms showing angular differences in PFD between rooms in S1–2diff (top) and S2–4diff (bottom). In both cases, distributions cluster significantly around 0°, confirming that HDCs preserved consistent tuning across rooms. Peak frequencies are indicated. V: V-test; Dir: mean PFD offset between rooms ± SD. ***p < 0.0001.

Similarly, in more complex environments, HDCs preserved this inter-room coherence. In S1-2diff and S2-4diff, vector lengths remained stable across rooms (**Supplementary Figure 6D**; S1-2diff, paired t-test: t(144) = 1.27, p = 0.21; S2-4diff, one-way ANOVA: F(2.383, 266.9) = 1.68, p = 0.18). Directional offsets again clustered near 0° (**Figure 4H**; S1-2diff, V-test for 0°: V = 106.37, p < 0.0001; S2-4diff, V-test for 0°: V = 280.97, p < 0.0001).

These findings demonstrate that HDCs maintain a stable directional signal across room boundaries and across environments. This contrasts with the realignment observed in MDCs, suggesting that HDCs encode an independent global reference frame.

### 3. Spatial coding in RSC extends beyond directionality

#### 3.1 Non-directional RSC cells exhibit room-specific spatial patterns aligned with environmental geometry

To determine whether the spatial coding of environmental structure in the RSC extends beyond directionally selective neurons, we next analyzed the activity of cells that did not meet the criteria for directional tuning. We investigated whether these non-directional neurons exhibited positional firing patterns that varied systematically across rooms, and whether any repetition or transformation of these patterns aligned with the geometry of the environment.

For each non-directional cell, we constructed spatial firing rate maps separately for each room and assessed the similarity of activity across rooms. To distinguish between patterns that are globally invariant and those structured according to local geometry, we computed two types of spatial correlation: (1) direct correlations of unrotated maps, and (2) correlations after rotating the maps to align salient structural features (e.g., doorways and visual cues; rotating through 180° in two-room environments and 90° in four-room environments). We considered three possible outcomes: (1) identical repetition across rooms, (2) repetition after rotation aligned with environmental geometry, and (3) distinct, non-repeating patterns (**Figure 5A,D**).

**Figure 5.**
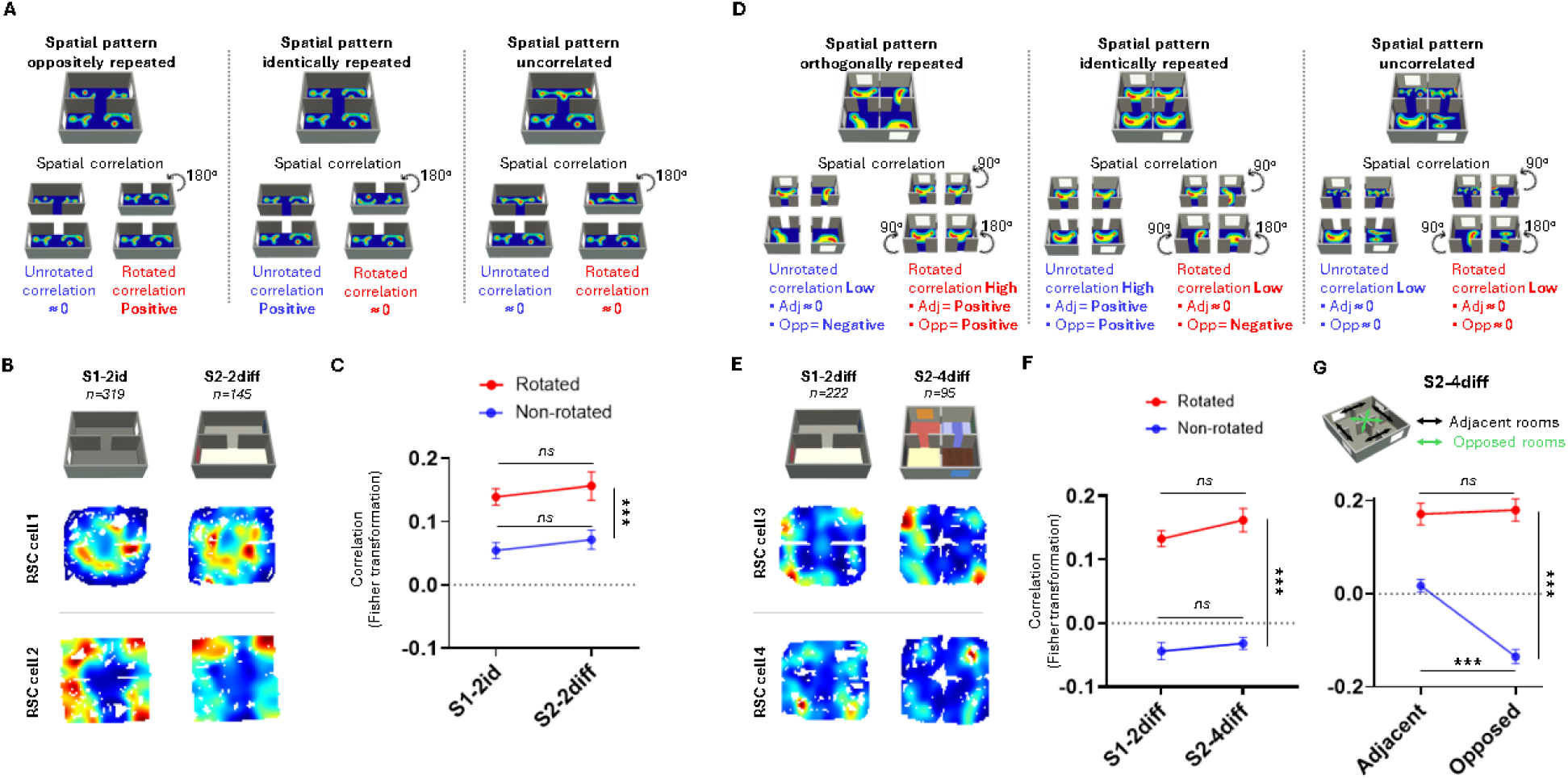
Non-directional RSC cells exhibit room-specific spatial patterns aligned with environmental geometry. **A.** Schematic of three hypothetical spatial firing patterns (top) and the corresponding spatial correlations (bottom) in two-room environments. **B.** Two examples of spatial ratemaps from non-directional RSC neurons recorded in two connected rooms that were either visually and tactilely identical (S1–2id) or distinct (S2–2diff). Both cells show oppositely repeated spatial patterns across rooms. **C.** Comparison of spatial correlations between unrotated and rotated ratemaps in S1–2id and S2–2diff. Rotated maps show significantly higher correlations than unrotated maps, regardless of room similarity. Dots represent individual data points; bars indicate mean ± SEM. Statistical comparisons were performed using a mixed ANOVA followed by Tukey’s post hoc test. **D.** Schematic of three hypothetical spatial firing patterns (top) and the corresponding predicted correlations (bottom) for unrotated and rotated comparisons between adjacent and opposite rooms. **E.** Two examples of spatial ratemaps from non-directional RSC neurons recorded in S1–2diff and S2–4diff environments. Cells exhibit oppositely aligned patterns in S1–2diff and orthogonally aligned patterns in S2–4diff. **F.** Comparison of spatial correlations between unrotated and rotated ratemaps in S1–2diff and S2–4diff. Rotated maps yield significantly higher correlations in both environments. Dots represent individual data points; bars indicate mean ± SEM. Statistical comparisons were performed using a mixed ANOVA followed by Tukey’s post hoc test. **G.** *Top*: Schematic representation of adjacent (black arrows) and opposite (green arrows) room comparisons in the 4-room environment. *Bottom*: Mean spatial correlations between adjacent and opposite room pairs. Unrotated maps show negative correlations between opposite rooms and near-zero correlations between adjacent rooms, whereas rotated maps yield positive correlations for both. Dots represent individual data points; bars indicate mean ± SEM. Statistical comparisons were performed using repeated-measures ANOVA followed by Fisher’s post hoc test. ***p < 0.0001.

First, we analyzed 319 cells recorded in a two-room environment with visually identical rooms (S1-2id) and then 145 of these cells in a structurally similar environment with visually and tactilely distinct rooms (S2-2diff). In both conditions, spatial correlations were significantly greater when maps were rotated, whereas unrotated maps showed values close to zero (**Figure 5B-C**). A mixed ANOVA revealed a significant effect of map alignment (F(3,462) = 11.45, p < 0.0001), but no effect of environment (F(1,462) = 0.09, p = 0.75), nor any interaction (F(3,462) = 0.07, p = 0.98). Post hoc tests confirmed that rotated maps consistently yielded higher spatial correlations (Tukey’s test, p < 0.0001).

We then performed the same analysis on 222 cells recorded in a distinct two-room environment (S1-2diff) and followed 95 of them in a distinct four-room environment (S2-4diff; **Figure 5E**). Again, spatial correlations were significantly higher for rotated maps than for unrotated ones (**Figure 5F**). A mixed ANOVA revealed a strong effect of rotation (F(3,630) = 47.99, p < 0.0001), with no environmental effect (F(1,630) = 0.03, p = 0.86) nor interaction (F(3,630) = 0.10, p = 0.96). Post hoc analyses confirmed higher correlations for rotated maps (Tukey’s tests, p < 0.0001).

These results show that non-directional RSC neurons encode room-specific spatial patterns that repeat across rooms following consistent geometric transformations (180° in two-room layouts and 90° in four-room layouts). These results also indicate that non-directional RSC neurons display spatial patterns that vary across rooms in a way that is consistent with the environmental structure.

#### 3.2 Room-to-room transformations reflect layout symmetry

To further examine how environmental geometry shapes these patterns, we next asked whether the transformations observed between room-specific maps depend on the spatial relationship between rooms. Specifically, we tested whether similarities in firing pattern differed between adjacent and opposite room pairs in the four-room environment (**S2-4diff**; **Figure 5G**), which are separated by angles of 90° and 180°, respectively. Based on the symmetry of the layout, we predicted that rotated correlations would remain high for both types of pair, while unrotated correlations would diverge, remaining near zero for adjacent rooms and becoming negative for opposite rooms.

To test this, we computed spatial correlations of firing rate maps for both adjacent and opposite room pairs, using both unrotated and appropriately rotated rate maps. A repeated-measures ANOVA revealed significant main effects of room relationship (adjacent vs opposite: F(1,94) = 21.29, p < 0.0001), map alignment (rotated vs unrotated: F(1,94) = 92.67, p < 0.0001), and a strong interaction between these two factors (F(1,94) = 30.25, p < 0.0001). Post hoc comparisons confirmed that unrotated correlations were significantly lower for opposite room pairs than for adjacent ones, consistent with a 180° spatial transformation (Fisher’s test, p < 0.0001). In contrast, rotated correlations were similarly high for both pair types, indicating that firing patterns were aligned once the correct transformation was applied (Fisher’s test, p = 0.68).

Animals tended to slow down near the doors as they are salient landmarks and strategic locations where animals often waited for the reward. We tested whether the doors and the behavioral patterns associated to them could account for the spatial activity of non-directional cells. We created population speed maps for successive sessions in two and four identical and different rooms environments (**Supplementary Figure 8A-B**) and we observed for all population speed maps a significant decrease of animals speed at the door compared to other part of the environment. Interestingly, we discovered that one-quarter to one-third of cells are positively or negatively modulated by running speed (**Supplementary Figure 9-10**). In contrast, similar analysis for population ratemaps of RSC cells revealed a homogeneous spatial distribution in both the two- and four-room environments and no changes of firing at the doors (**Supplementary Figure 8C-D**). These results indicate that neither the salience of the doors nor local variations in running speed can explain the spatial activity patterns observed between connected rooms. These results suggest that RSC non-directional cells reflect the geometric segmentation of the environment rather than specific local features or stereotype movement within it.

Altogether, these findings demonstrate that spatial representations in the RSC are both room-anchored and organized according to predictable geometric relationships between rooms. The ability to recover strong correlations after appropriate rotation indicates that spatial codes incorporate both local reference frames and global structure of the environment, supporting a flexible, geometry-aligned encoding system of complex environments.

### 4. Divergent spatial coding in RSC and hippocampus

#### 4.1 RSC spatial codes are stable over time

To determine whether the spatial coding observed in the RSC is stable over time, we tracked non-directional RSC neurons as animals were repeatedly exposed to multi-room environments. We recorded from 40 RSC neurons with a high spatial information content (≥0.9, see spatial information distribution on **Supplementary figure 7**) during a four-session protocol alternating between visually identical and distinct two-room configurations (S1-2id, S2-2diff, S3-2id, and S4-2diff). Figure 6A shows example cells that illustrate how the structured, oppositely aligned spatial patterns observed between rooms were preserved across sessions, despite the alternating sensory conditions.

**Figure 6.**
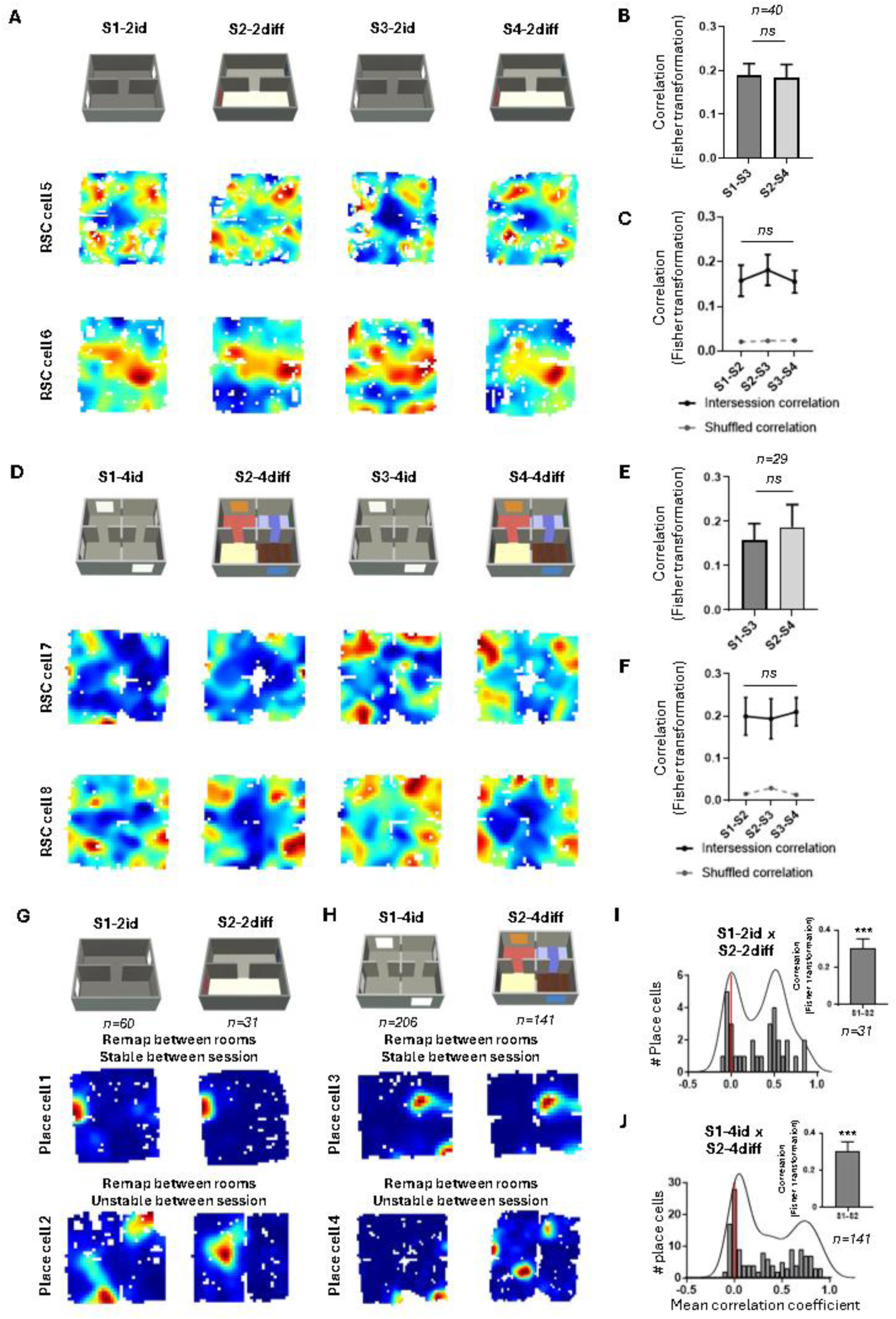
Divergent spatial coding in RSC and hippocampus. A. Two examples of spatial ratemaps from non-directional RSC neurons recorded across four consecutive sessions alternating between two environments composed of two connected rooms that were either visually and tactilely identical (S1-2id, S3-2id) or distinct (S2-2diff, S4-2diff). Only RSC neurons with a high spatial information content (≥0.9) are preserved for this analysis. They show oppositely repeated spatial patterns that are preserved across sessions. **B.** Mean spatial correlation values for repeated environments (S1–2id × S3–2id and S2–2diff × S4–2diff). No significant differences were observed between the two conditions. Bars represent mean ± SEM. Statistical comparison was performed using a paired t-test. **C.** Mean spatial correlations across consecutive sessions in the full protocol (S1–2id × S2–2diff, S2–2diff × S3–2id, S3–2id × S4–2diff). Correlations remained positive and significantly higher than shuffled controls. Bars represent mean ± SEM. Statistical comparison was performed using one-way ANOVA. **D.** Two examples of spatial ratemaps from non-directional RSC neurons recorded across four successive sessions alternating between two environments composed of four connected rooms that were either visually and tactilely identical (S1–4id, S3–4id) or distinct (S2–4diff, S4–4diff). Cells exhibit orthogonally aligned patterns that remain consistent across sessions. **E.** Mean spatial correlation values for repeated environments (S1–4id × S3–4id and S2–4diff × S4–4diff). No significant differences were observed between the two conditions. Bars represent mean ± SEM. Statistical comparison was performed using a paired t-test. **F.** Mean spatial correlations across successive sessions in the 4-room protocol. Correlations remained positive and significantly higher than shuffled controls. Bars represent mean ± SEM; statistical comparison was performed using one-way ANOVA. **G.** Examples of hippocampal place cells recorded in S1–2id and S2–2diff. Cells show either repeated or remapped firing fields across rooms, with most exhibiting remapping. Opposite firing fields were never observed. **H.** Examples of hippocampal place cells recorded in S1–4id and S2–4diff. Place cells almost exclusively remap across rooms, with no evidence of orthogonally repeated fields. **I.** *Left:* Distribution of spatial correlation values between S1–2id and S2–2diff for all place cells, overlaid with kernel density estimates. The bimodal distribution indicates the coexistence of stable and unstable spatial activity across sessions. *Right:* Mean spatial correlation values between S1-2id and S2-2diff. Bars represent mean ± SEM. Statistical comparison was performed using a one sample t-test against 0. ***p < 0.0001. **J.** Same as in I, but for the four-room condition (S1–4id × S2–4diff). The distribution similarly reveals clusters of stable and unstable place cell activity. Bars represent mean ± SEM. Statistical comparison was performed using a one sample t-test against 0. ***p < 0.0001.

Quantitative analyses confirmed this visual impression. Spatial correlations between matching sessions were significantly greater than zero (one-sample t-tests against 0: all p < 0.0001) and did not differ between visually identical and distinct environments (**Figure 6B**: paired t-test: t(39) = 0.16, p = 0.87). Furthermore, correlations across all successive session pairs were consistently positive, significantly above zero (one-sample t-test against 0: p < 0.0001 for all comparisons), greater than shuffled controls (95th percentile thresholds) and exhibited no significant differences between session pairs (**Figure 6C**: one-way ANOVA, F(1.90, 74.08) = 0.22, p = 0.79), supporting the conclusion that spatial activity remained stable throughout the recording protocol.

To assess whether this temporal stability generalized to more complex layouts, we conducted a parallel experiment using four-room environments (S1-4id, S2-4diff, S3-4id, and S4-4diff; **Figure 6D**). We followed 29 RSC neurons across sessions and again observed preserved spatial transformations across rooms, including consistent orthogonal patterning. Correlations between repeated environments were significantly greater than zero (one-sample t-test against 0: p < 0.001 for both) and did not differ between layout types (**Figure 6E**: paired t-test: t(28) = 0.62, p = 0.54). Spatial correlations remained positive across all successive sessions, significantly above zero (one-sample t-test against 0: p < 0.0001 for all comparisons), greater than shuffled controls (95th percentile thresholds) and exhibited no change across the four-session protocol (**Figure 6F**: one-way ANOVA, F(1.83, 51.12) = 0.08, p = 0.91).

Together, these results demonstrate that non-directional RSC neurons maintain stable, structured spatial representations over time, despite perceptual changes. This temporal consistency strengthens the hypothesis that the RSC supports a robust spatial code anchored to room structure and layout.

#### 4.2 Hippocampal place cells remap between rooms

In light of the geometry-aligned spatial transformations observed in the RSC, we investigated whether the hippocampus exhibits a comparable organizational structure or whether it relies on distinct principles. Although it is known that hippocampal place cells either repeat or remap across connected rooms (Cheng *et al*., 2024; Duvelle *et al*., 2021; Grieves *et al*., 2016; Paz-Villagrán *et al*., 2004; Skaggs & McNaughton, 1998; Spiers *et al*., 2015), it is unclear whether their activity can show structured transformations similar to those observed in the RSC. Specifically, we tested whether place cells display repeated firing fields aligned with room geometry, or whether their representations change independently across connected rooms.

To address this, we recorded the activity of hippocampal neurons in 4 animals (**Supplementary Figure 11**) as they freely moved in both two-room and four-room environments. We then compared neurons spatial firing patterns across rooms and across sessions. We systematically evaluated whether spatial correlations reflected geometric transformations, identical repetition, or complete remapping, using both unrotated and rotated spatial rate maps to detect possible alignment effects (**Figure 5A,D**).

We recorded 57 place cells in the two visually identical rooms environment (S1-2id) and tracked 31 of these place cells in the structurally similar, perceptually distinct environment (S2-2diff; **Figure 6G**). Cells with place fields located at the door were excluded from analysis to avoid spatial ambiguity (**Supplementary figure 12)**. Using a correlation threshold of 0.4 to define significant spatial similarity (as in Spiers *et al*., 2015), we found that the vast majority of place cells remapped between rooms in both environments (91%), with only a small subset exhibiting repeated fields (9%), and none showing the opposite or orthogonally repeated activity patterns as observed in the RSC.

We then analyzed hippocampal activity in four-room environments. We recorded 206 place cells in S1-4id and followed 141 of them in the perceptually distinct S2-4diff condition (**Figure 6H**). Remapping between rooms was almost universal in both conditions (99.5% and 100%, respectively), and, once again, no orthogonally repeated patterns were detected.

Overall, these results demonstrate that, unlike the RSC, the hippocampus does not generalize spatial codes across rooms in a geometry-dependent manner. Rather, its representations reflect room-specific remapping with no evidence of consistent alignment to environmental geometry.

#### 4.3 Place cell activity is stable across sessions, despite remapping

We next assessed whether hippocampal place fields were stable across sessions. Although RSC non-directional spatial activity is stable across sessions, hippocampal place cells could show either stable or non-stable place field activity. In the two-room environments, the correlation between S1-2id and S2-2diff followed a bimodal distribution (**Figure 6I**: kernel peaks at 0 and 0.55), which suggests that there is variability in the stability of individual place cells that depends on the session (**Supplementary Figure 13**). Nevertheless, overall spatial correlations were significantly greater than zero (one-sample t-test against 0: t(30) = 5.63, p < 0.0001), indicating a degree of temporal consistency despite spatial reorganization.

A similar pattern was seen in the four-room environment. Correlations between S1-4id and S2-4diff also followed a bimodal distribution (**Figure 6J**: peaks at 0.05 and 0.7), again suggesting session-dependent variability in field stability. Nevertheless, mean correlations were also significantly above zero (one-sample t-test against 0: t(140) = 10.39, p < 0.0001), reflecting a consistent, albeit locally variable, session-level structure.

Together, these results show that while hippocampal place cells remap between rooms during a single session, their spatial activity can remain stable across sessions.

## Discussion

Navigating the real world involves interacting with environments that are made up of separated but interconnected spaces. The discontinuities imposed by the structural boundaries raise the question of how the brain constructs coherent internal representations across segmented spaces. While previous studies have demonstrated the contribution of the retrosplenial cortex (RSC) to spatial orientation (Vann *et al*. 2009; Alexander *et al*., 2023), our findings expand upon this by revealing its role in encoding structured representations that align with both environmental geometry and room boundaries. By combining recordings from directionally tuned and non-directional neurons, we reveal that the RSC supports a multi-scale spatial code: head direction cells provide a stable global reference, whereas multidirectional and non-directional cells exhibit room-specific firing patterns organized according to the environmental layout. In contrast, hippocampal place cells remap between rooms without preserving geometric alignment, highlighting a dissociation in spatial coding. Overall, these findings establish the RSC as a pivotal integrative hub for spatial segmentation in complex environments, where complementary neural populations converge to support a geometry-aligned internal map.

A central component of the RSC’s spatial code is its organization of directional tuning across rooms. Our results reveal the coexistence of two distinct yet complementary populations: head direction cells (HDCs), which maintain a stable preferred firing direction across rooms, and multidirectional cells (MDCs), which shift their preferred firing directions in a room-specific manner. This dual coding scheme suggests that the RSC encodes direction across multiple spatial scales, flexibly linking local frames to a unified global orientation -a structure that is well suited to navigate in segmented environments.

Crucially, our findings show that MDC tuning reflects environmental structure rather than perceptual ambiguity. Previous studies have identified MDCs in visually identical rooms that were distinguished only by odor cues (Jacob *et al*., 2017; Zhang *et al*., 2022), leaving open the possibility that directional shifts are driven by confusion in visual sensory cues. To address this, we recorded in environments where rooms were clearly differentiated by visual and tactile features; and observed that MDCs still exhibited geometry-aligned tuning. Their preferred directions were reversed by 180° in two-room layouts and rotated by 90° in four-room configurations, which is consistent with the spatial arrangement of the rooms. One might hypothesize that specific environmental cues, such as door orientation or landmark positioning, drive these tuning shifts. However, our results argue against this interpretation. In four-room environments, each room had multiple entry points, yet MDCs never exhibited multidirectional activity within a single room. Furthermore, previous studies has shown that MDC tuning persists in complete darkness (Jacob *et al*., 2017; Zhang *et al*., 2022), indicating that visual cues are not required.

Altogether, these findings suggest that MDCs are not anchored to one specific local sensory feature, but rather reflect the geometric segmentation of space into discrete rooms. This tuning pattern is consistent with the idea that MDCs are anchored to local reference frames defined by spatial boundaries. Each walled room provides a distinct directional context that integrates spatial and non-spatial features (Jacob *et al*., 2017; Zhang *et al*., 2022; Marchette *et al*., 2014). In contrast, HDCs maintain a global directional anchor, remaining stable across rooms regardless of their geometry. The coexistence of these two systems may enable the RSC to relate distinct local frames *via* a stable global reference, thereby supporting an internal model of spatial topology, i.e. the relative orientation and connectivity of discrete rooms. In undivided environments, local and global frames may converge into a single reference frame, which could explain why multidirectional tuning disappears or switches to unidirectional tuning in open fields (Jacob *et al*., 2017; Zhang *et al*., 2022).

Beyond their complementary coding roles, MDCs and HDCs exhibit fundamentally distinct properties in different environmental configurations, suggesting that they rely on different neural mechanisms. MDCs consistently showed directional shifts aligned to room geometry while maintaining unipolar tuning within each room. This suggests that their multidirectionality arises from a sequence of room-specific directional anchors. In contrast, HDCs preserved a stable preferred direction across all environments and never adopted multidirectional patterns. This dissociation challenges the concept of a unified head direction system governed by a single attractor network (Zhang, 1996; Redish *et al*., 1996; Clark & Taube, 2012). Instead, our findings support the existence of distinct directional subsystems that possibly rely on parallel attractor-like dynamics or alternative mechanisms (Page & Jeffery, 2018; Samsonovich, 2018; Kornienko *et al*., 2018). In particular, Page and Jeffery (2018) proposed that RSC neurons could integrate global HD signals with local landmarks, enabling bidirectional tuning to emerge *via* Hebbian learning. While originally framed around symmetry-induced bidirectionality, this model provides a compelling explanation for our findings: HDCs may reflect a stable, global attractor, whereas MDCs flexibly bind to local frames through experience-dependent associations.

This dual system also offers a potential explanation for the discrepancies observed in human neuroimaging studies. These studies have alternately implicated the RSC in either local (Marchette *et al*., 2014; Shine *et al*., 2016) or global (Kim & Maguire, 2018; Peer *et al*., 2019; Sulpizio *et al*., 2013) spatial reference frames. Our results suggest that the RSC supports both types of reference frame, enabling a flexible integration of room-specific orientation and global spatial coherence in complex environments.

While HDCs and MDCs account for only a small proportion of the RSC population, most neurons do not exhibit clear directional tuning. Previous studies have shown that these non-directional neurons can encode spatial position, movement, or contextual variables (Alexander *et al*., 2015, 2017, 2020; Keshavarzi *et al*., 2022; Mao *et al*., 2017, 2018; Miller *et al*., 2019, 2021; Vedder *et al*., 2016; Xie *et al*., 2014). Our findings extend this view by showing that non-directional RSC neurons also exhibit organized spatial activity that aligns with environmental geometry. Specifically, their firing patterns were mirror-symmetric across two rooms and rotated by 90° across four rooms, consistent with the global layout. These transformations are similar to those observed in MDCs, suggesting that geometry-aligned segmentation is not limited to directional neurons, but rather represents a more general organizational principle within the RSC. By encoding both local reference frames and their spatial transformations, the RSC can create a flexible, geometry-based map of complex environments. This map not only captures the content of individual rooms but also their spatial relationships, providing a framework for large-scale spatial reasoning.

A key question is whether this spatial segmentation also reflects contextual encoding. In contrast to previous findings linking RSC activity to contextual remapping and episodic memory (Alexander *et al*., 2015, 2023; Mao *et al*., 2017; Miller *et al*., 2021), we observed here that non-directional cells exhibited similar representations across connected rooms that were specifically aligned with the geometry of each room. If rooms are considered not only as geometric units, but also as distinct functional contexts, defined by local cues, task demands, or behavioral contingencies, then this geometry-aligned spatial segmentation could serve as a proxy for contextual boundaries. By anchoring neural activity to room-specific frames, the RSC could facilitate transitions between spatial contexts and support the organization of episodic memory. Thus, spatial and contextual segmentation may thus arise from a shared computational principle: the partitioning of continuous experience into discrete, navigation-relevant episodes.

This interpretation is reinforced by the contrasting coding scheme observed in the hippocampus. While the RSC maintained geometry-aligned representations across rooms, hippocampal place cells adopt a markedly different coding strategy. In both two- and four-room environments, they consistently remapped between rooms, generating distinct spatial firing fields without preserving any regular geometric transformation. This divergence highlights that each brain region employs fundamentally distinct coding rules.

Remapping is widely regarded as a hallmark of contextual differentiation, with place cells generating unique representations for environments perceived as distinct in structure, content, or behavioral relevance (Muller & Kubie, 1987; Anderson & Jeffery, 2003; Leutgeb *et al*., 2005). In multicompartment settings, hippocampal cells typically adopt one of two mutually exclusive codes: they either repeat their firing fields across similar rooms, or they remap completely (Grieves *et al*., 2016, 2017; Harland *et al*., 2017; Paz-Villagrán *et al*., 2004; Spiers *et al*., 2015). This mutual exclusivity suggests that the hippocampal code commits to a single spatial framework at a time: either generalizing across rooms, or differentiating them based on contextual boundaries. The consistent remapping observed in our study suggests that each room was encoded as a distinct contextual entity, even when rooms shared a common geometry.

Previous studies have shown that the geometric layout of rooms within a multicompartment environment can determine whether place cell activity repeats or remaps across rooms. Specifically, place cells tend to display repeated firing fields in environments with parallel connected rooms, whereas in radial configurations, they remap between rooms (Grieves *et al*., 2016). The repetition of firing fields in parallel rooms emphasizes that place cell activity is anchored to the local room scale. On this basis, we can hypothesize that when rooms have different orientations, place cells remap, coding each room separately rather than exhibiting a global activity with multiple fields. Consistent with this, our experiment have rooms arranged in opposing and orthogonal orientations and place cells predominantly exhibited remapping across rooms. Moreover, the directional signal appears to be a key factor in shaping place cell representations in connected environments. Harland *et al*. (2017) showed that lesions of the lateral mammillary nuclei — which disrupt the directional signal — resulted in an increased repetition of place cell firing fields when animals are exposed to radial-rooms configuration. As the MDCs signal varies across connected rooms with different orientations, it may provide orientation-specific input to the hippocampus, leading to place cells remapping. Recording MDC activity in environments with parallel connected rooms would be informative, as the shared orientation between rooms may result in a uniform directional signal, which could support repeated coding by place cells.

In summary, our findings demonstrate that the RSC encodes a structured representation of complex environments by aligning local and global reference frames. This multi-scale coding strategy enables the RSC to support the simultaneous representation of discrete spatial units and their global organization. In contrast, the hippocampus overlays this scaffold with context-specific representations, enabling flexible encoding of experience and memory. This division may support flexible navigation, by organizing spatial information across multiple spatial levels.

## Methods

### Subjects

A total of 15 adult Long Evans rats (Charles River Laboratories), 12 males and 3 females weighing between 250 and 350g were housed in individual cages (40 cm long x 26 cm wide x 16 cm high) with *ad lib* food and water and kept in a temperature-controlled room (20°C +/- 2°C) with natural light/dark cycle. One week after arrival, animals were handled daily by the experimenter for 7 days. During this period, they were habituated to the recording apparatus by letting them explore the environment for 10–20 min per day.

All experimental procedures were conducted with the approval of the local French National Ethics Committees and in accordance with the EEC (2010/63/UE) guidelines for the care and use of laboratory animals. The present study was specifically approved by Neurosciences Ethics Committee N°71 of the French National Committee of animal experimentation.

### Microdrives and surgery

Recordings were made using tetrodes, each composed of four twisted 25 μm polyimide-coated platinum-iridium (90%/10%) wires (California Fine Wire, CA), attached to an Axona microdrive (Axona Ltd, St Albans, UK). Surgery was performed under sterile conditions and general anesthesia (Isoflurane, 4% for induction, 2% to 0.5% for maintenance), an opioid analgesic (Buprenorphine, 0.05 mg/kg, SC) is administered at least 30 minutes before. Tetrodes were then stereotaxically implanted in the dysgranular RSC (n = 14 rats; coordinates in mm from Bregma: AP: -5.0 to -5.5mm, ML: +/-0.75 to 1.0mm and DV: 0.4 to 0.5mm under the dura) and/or the dorsal hippocampus (n = 4 rats; coordinates in mm from Bregma: AP: -3.8, ML: +/-3 and DV: 1.5mm under the dura) either in the left or right hemisphere. As a postoperative treatment, rats received an injection of antibiotic (Oxytetracycline, 10 mg/kg) and of anti-inflammatory/analgesic (Carprofen, 5 mg/kg). After surgery, the rats were allowed 7 days of recovery before the experiment began. They were then placed on a food deprivation schedule that kept them at 90% of their free-feeding body weight.

### Apparatus

The 2- and 4-connected-rooms environments were derived from a 150 x 150 cm square arena with 60 cm high grey walls and covered with a gray vinyl sheet that can be wiped with a moistened sponge to neutralize smells. The apparatus was isolated from the rest of the room by an opaque circular curtain 260 cm in diameter. Ceiling lights lighted the environments homogeneously and a radio fixed to the ceiling above the environments was used to mask uncontrolled directional sounds. The unit recording system and equipment for controlling the experiment were in an adjacent room.

To create a two visually identical opposite connected rooms, we divided the arena in 2 equal rectangular rooms (75 x 150 cm) by inserting a 60 cm high wall with a door at the center (an aperture of 10 cm at the base and 15 cm at the top) (**Supplementary figure 1A**). For the four visually identical orthogonal connected rooms, we divided the arena in four equal square rooms (75 x 75 cm) by inserting two 60 cm high walls perpendicularly, each having two doors allowing to connect each room with the two adjacent ones (**Supplementary figure 1B**). We polarized each room with identical white cue cards (30 x 40 cm, one per room) attached to the opposite short walls in the 2-identical-rooms environment, or to the orthogonally arranged walls in the 4-identical-rooms environment. To maximize the difference between the rooms, in the 2- and 4-different-rooms environments, each cue card had a specific pattern (a circle, stripes, a square, a star, a triangle) and each floor was covered with vinyl of different texture and visual pattern (**Supplementary figure 1C and D**).

### Recording setup

Both screening and recordings were performed in the experimental room. A counterbalanced cable was attached at one end to an automatic commutator and at the other end to the headstage containing a field effect transistor amplifier for each tetrode wire. The signals from each tetrode wire were amplified 10 000 times, bandpass filtered between 0.3 and 3 kHz with Neuralynx amplifiers (Neuralynx, Bozeman, MT, USA), digitized (32 kHz), and stored by DataWave Sciworks acquisition system (DataWave Technologies, Longmont, CO, USA). Two light-emitting diodes (LED), one red and one green attached to the headstage provided the position and the orientation of the rat’s head. The LEDs were imaged with a CCD camera fixed to the ceiling above the maze, and their position was tracked at 25 Hz with a digital spot-follower. A food dispenser of 20 mg food pellets was located 2 m above the environment. Animals were moving freely while food was randomly dropped on the floor to motivate them to explore the entirety of the environment.

### Recording protocols

Recordings began one week after the surgery. The animals were daily screened in an individual box to identify single unit activity. When no single unit was observed or after a completed recording protocol, the tetrodes were lowered 50 μm. When one or several units were isolated, we started one of the 3 recording protocols of the study:

1. one session in the 2-identical-connected-rooms environment followed by one session in the 2-different-connected-rooms environment. This recording protocol could be repeated twice in succession in order to assess the spatial stability of non-directional neurons in the RSC.
2. one session in the 2-different-connected-rooms environment followed by one session in the 4-different-connected-rooms environment.
3. one session in the 4-identical-connected-rooms environment followed by one session in the 4-different-connected-rooms environment. This recording protocol could be repeated twice in succession in order to assess the spatial stability of non-directional neurons in the RSC.

Each session lasted 10-20 minutes, and the floor was cleaned between sessions at the end of the protocol. Between successive sessions, the rats were not disconnected but were removed from the apparatus and placed in a random location in an individual box to let the experimenter manipulate the apparatus (changing the visual cues, the floors and/or the central walls, cleaning the walls and the floor). Animals were then mildly disoriented by rotating the box before the next session.

### Data analyses

All data analyses (except for the spike sorting) were performed with Matlab® custom scripts.

#### Spike sorting

Spike sorting was performed manually using the graphical cluster-cutting software Offline Sorter (Plexon). Units selected for analyses had to be well-discriminated clusters with spiking activity clearly dissociated from background noise. Units having inter-spike intervals < 2 ms (refractory period), were removed due to poor isolation. To prevent repeated recordings of the same cell over days, clusters that recurred on the same tetrodes in the same cluster space across recording sessions were only analyzed one time.

#### Directionality analyses

The rat’s head direction was calculated for each tracker sample from the projection of the relative position of the two LEDs onto the horizontal plane. The directional tuning for each cell was obtained by dividing the number of spikes when the rat faced a particular direction (in bins of 6°) by the total amount of time the rat spent facing that direction. The cell’s PFD is the circular mean of the directional tuning. Further analyses were only performed on cells having a peak firing rate exceeding 1 Hz.

#### Direction cells selection

To assess the directional specificity of **HDCs,** the Rayleigh vector length was used, with significant directionality being assigned to cells whose Rayleigh vector score exceeded 0.2. **Bidirectional cells** (**BDCs)** were identified with a flip score exceeding 0.6, as in a previous study in which it was necessary to select BDCs (*Jacob et al. 2017*). The flip score was calculated with an autocorrelation procedure, by rotating the polar firing rate plot in steps of 6° and calculating the correlation between the rotated and unrotated plots at each step. The flip score for each cell was defined as the difference between the correlations at the expected peak (180°) and the expected trough (90°). Note that, unlike Jacob et al. (2017), who identified ‘within-compartment bi-directional cells’ that fired bidirectionally in both rooms of the environment and ‘between-compartment bi-directional cells’ that changed their preferred firing direction between rooms, we did not observe these two subpopulations of bi-directional cells. **MDCs** were defined as cells showing multidirectional activity in the 4-connected-rooms environment but showing unipolar activity in individual rooms. MDCs were selected based on one criterion: the Rayleigh vector score calculated from the directional activity in each of the four rooms had to exceed 0.2 in at least one of the four rooms.

#### Comparison of the BDCs and HDCs activities between environments

We analyzed whether cells changed their activity between 2-identical and 2-different-connected-rooms environments by comparing the peak firing rate and either the flip scores or the vector lengths, respectively for BDCs and HDCS, using a repeated measure ANOVA.

#### Characterization of the MDCs activity

Since the rooms in the 2-connected-rooms environments are oppositely arranged, we tested whether the PFDs were separated by 180° between rooms. For each cell, we calculated the differences between the PFDs of the two rooms and tested whether they were distributed around 180°, by extracting the circular mean value with its corresponding angle and the circular V-test value for 180°. Similarly, in the 4-connected-rooms environment that are orthogonally arranged, we tested whether the PFDs were separated by 90° between rooms. We calculated the differences between the PFDs of adjacent rooms and tested whether they were distributed around 90°, by extracting the circular mean value with its corresponding angle and the circular V-test value for 90°.

For HDCs we carried out similar analyses in the 2- and 4-connected-rooms environments, assuming similar PFDs between rooms. We tested whether the PFDs were separated by 0° between room by calculating the differences between the PFDs of adjacent rooms and tested whether they were distributed around 0°, by extracting the circular mean value with its corresponding angle and the circular V-test value for 0°.

#### Spatial ratemaps and place cell selection

Spatial ratemaps were generated by dividing the environment into square bins, each 2.5 × 2.5 cm in size. Spikes per bin were divided by the time spent in that bin to provide a raw firing ratemap. Smoothed firing ratemaps were then generated using a boxcar procedure in which the firing rate in each bin was replaced by that of the mean of itself plus the immediately surrounding bins. The firing rate of the cell was color-coded from low (light blue) to high (dark red). In both raw and smoothed ratemaps, pixels that were not visited by the rat were displayed in white.

For each map, we calculated two classical measures of spatial selectivity:

– the spatial coherence is calculated on the non-smoothed (raw?) rate map as the spatial autocorrelation of the non-smoothed place field and measures the extent to which the firing rate in a particular bin is predicted by the average rate of the eight surrounding bins.
– the spatial information content (Skaggs *et al*., 1996), expressed in bits per second and is calculated as follows:

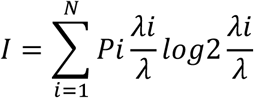

Where 𝑃(𝑖) is the occupancy probability of bin, 𝜆𝑖 is the mean firing rate in bin and 𝜆 is the overall mean firing rate.

Hippocampal cells were categorized as place cells if they satisfied 4 criteria in at least one room of the 2- or 4-connected-rooms environments:

1– the place field should include at least nine contiguous pixels of the non-smoothed ratemap with a firing rate above the mean firing rate (averaged over the environment).
2– the peak firing rate should exceed 1Hz.
3– the spatial coherence score should exceed 0.4.
4– the spatial information content should exceed 1.

#### Spatial correlation

To analyze whether both place cells and RSC neurons have repeated, opposite or unrelated spatial activity between rooms, we constructed for each cell the ratemaps of each separated room. For the 2-connected-rooms environments, we calculated the Pearson product-moment correlation coefficient on Fisher-transformed values between the 2 non-smoothed spatial ratemaps either unchanged or rotated by 180°, to match the position of the door and the visual cue cards. For the 4-connected-rooms environments, we calculated all possible spatial correlations between any two rooms (4 correlations for adjacent rooms and 2 correlations for opposite rooms) which we either averaged altogether, or averaged according to whether the rooms were adjacent or opposite. In order to assess whether place fields rotated or repeated across rooms, we calculated the mean spatial correlation of the unchanged rooms and after rotating three of the four rooms by 90°, -90° and 180° in order to overlap the position of the doors and the visual cue cards.

Finally, to test the stability of RSC and place cells spatial activity across our multicompartment environments composed of identical and different rooms we calculate the spatial correlation between successive sessions of 2-identical and 2-different-connected-rooms environments. We focused on RSC cells exhibiting spatial coding by selecting those with a spatial information content greater than 0.9 (a threshold at which the distribution of spatial information content of our RSC cells shows a clear demarcation; **Supplementary Figure 7**) along with all recorded place cells. In the same way we calculated the spatial correlation between successive 4-identical and 4-different-connected-rooms environments. The intersession correlations in the 2- and 4-rooms environment were compared to a bootstrap distribution consisting of correlating each cell ratemap with a randomly selected cells (**repeated 1000 times**).

#### Population speed and rate maps

To analyze whether doors influence RSC spatial activity, we analyzed animal speed for all sessions having RSC neurons recordings and population rate maps for all non-directional RSC cells. Population speed map : for each recording, we calculated for each sample position the distance that we divided by the tracking sampling frequency to extract instantaneous speed, then we created a spatial speed map by dividing the environment into square bins, each 2.5 × 2.5 cm in size and assigning in each bin the average speed. Finally, we average all speed maps for all sessions in the two rooms environment (35 recordings) and in the four-rooms environment (19 recording sessions). Population rate map : we used the spatial ratemap methods described above for RSC non-directional neurons that were recorded during the 4 successive sessions (session 1 with identical rooms, session 2 with different rooms, session 3 with identical rooms and session 4 with different rooms) either in the two-rooms environments (222 neurons) and four-rooms environments (76 neurons). Finally, we averaged all ratemaps for each session. We assessed statistical significance at the door position by extracting from each population maps either average speed or average firing rate of the nine contiguous bins centered to the doors (1 door for the two-rooms and four doors for the four-rooms environments), that we compared to a shuffled distribution. This shuffling were made for each session by extracting 1 000 times the speed or the firing rate in nine contiguous bins positioned randomly in the population map, including the position of the doors. Then we extracted the 5^th^ percentile of the shuffling speed or firing rate values as significant thresholds. A speed value or a firing rate at doors below these thresholds indicate a significant change at the doors.

### Statistical analyses

Statistics were performed with GraphPad Prism 9® and included Mann-Whitney U test, one-sample Wilcoxon test, one-sample Student’s t-test, two-samples Student’s t-test either paired or unpaired, one-way ANOVA, two-way ANOVA and repeated-measure ANOVA, Tukey’s post hoc, were performed when allowed. Rayleigh tests for uniformity and circular V-tests were performed with Matlab®.

### Histology

At the completion of the experiment, after induction of deep sedation (Domitor, 500μg/kg) rats received an overdose of barbiturate (Euthasol, 200 mg/kg). The brains were removed and frozen with isopentane cooled at -80°C. Then 25 µm-thick coronal sections were mounted on glass slides (SuperFrost Plus®, ThermoFisher) and stained with thionine. The position of the tips of the electrodes was determined from digital pictures, acquired with a Leica Microscope (Wetzlar, Germany), and imported in an image manipulation program (Gimp 2.8, distributed under General Public License). Delimitations of the RSC and hippocampus were based on Paxinos and Watson’s stereotaxic atlas (*Paxinos & Watson, 2007*), recording sites were determined by measuring backwards from the deepest point of the track.

## Acknowledgments

This work was supported by a grant from the Agence Nationale de la Recherche (ANR-21-CE37-0006-01), by the Centre National de la Recherche Scientifique (CNRS) and Aix-Marseille University (AMU). We thank the Spatial Cognition group for their insightful comments on the manuscript, S. Quitard for the histological work, the Olympique de Marseille and Jul. C. Laurent salary for this work was provided by the ANR grant.

## Author contributions

C.L., P.Y.J. and F.S. designed the study. C.L. and P.Y.J. performed surgeries and recordings. C.L. & P.Y.J. analyzed data. All authors interpreted data and discussed results. C.L., N.E.M and P.Y.J. wrote the manuscript. All authors commented and edited the manuscript.

