## Supplemental figures for "Divergent spatial codes in retrosplenial cortex and hippocampus support multi-scale representation of complex environments"

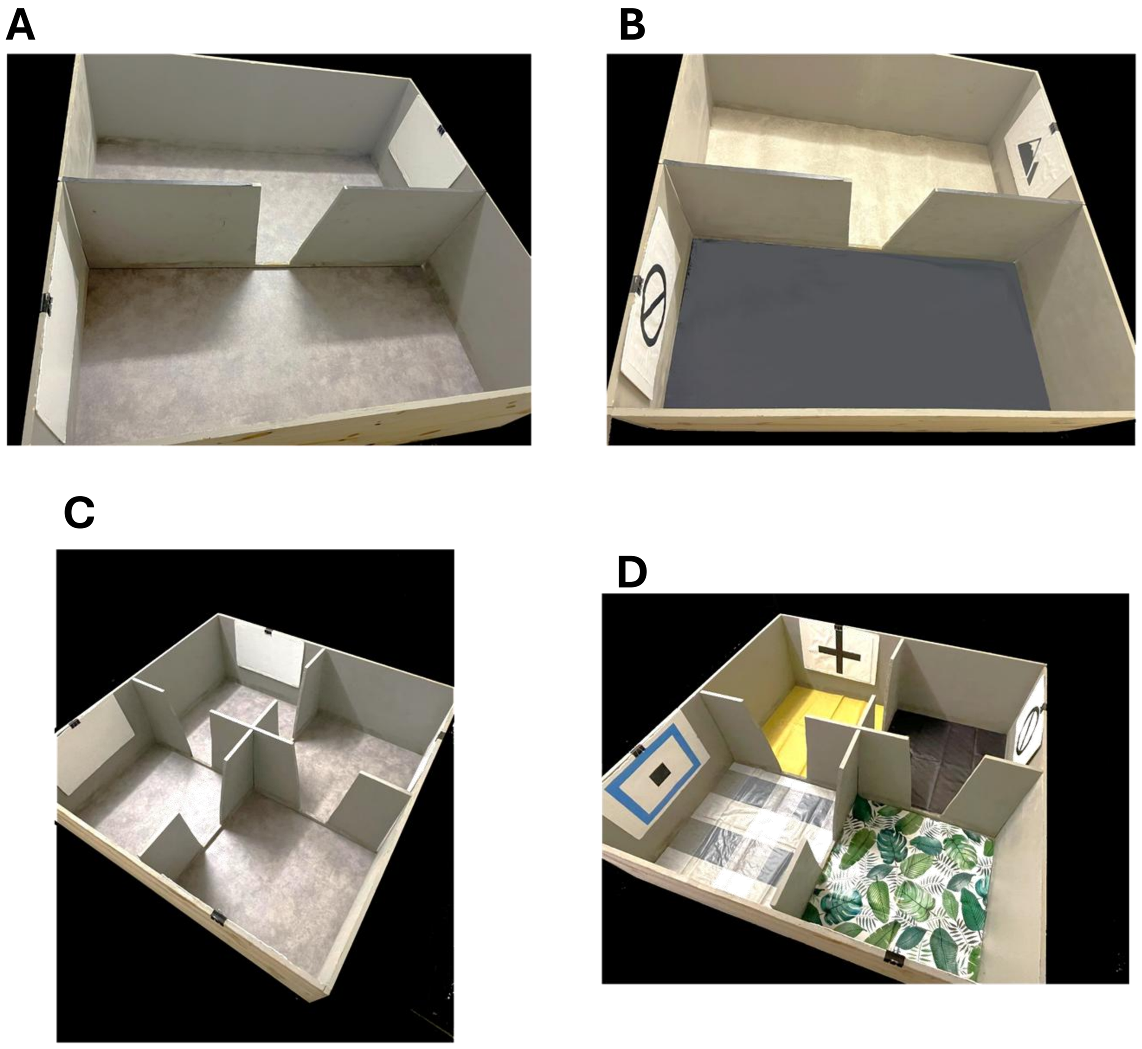

**Supplementary figure 1 – Pictures of the 4 connected rooms environment.**
**A.** The 2 identical connected rooms environment (same visual cues and floor). **B.** The 2 different connected rooms environment (different visual cues and floor colors and textures). **C.** The 4 identical connected rooms environment (same visual cues and floor). **D.** The 4 different connected rooms environment (different visual cues and floor colors and textures).

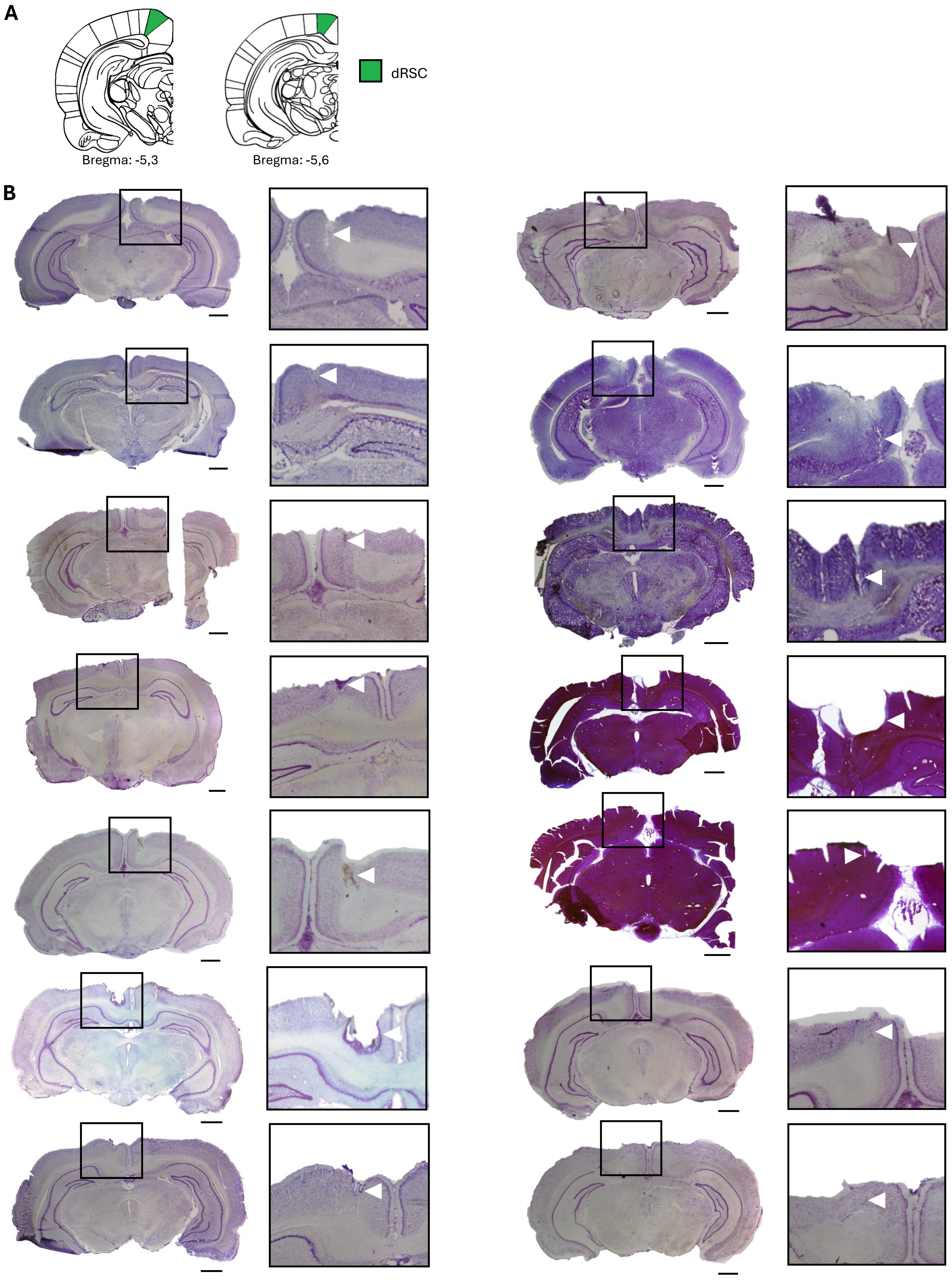


**Supplementary figure 2 – Histology retrosplenial cortex.**

**A.** Diagram showing the location of the dysgranular retrosplenial animals (RSC) adapted from Paxinos G, Watson C. The Rat Brain in Stereotaxic Coordinates. 4th Edition. Academic Press 1998. **B.** Coronal slices through the region of electrode implantation from the RSC (black insets) with scale bars indicating 1 mm. Bottom right: magnification of insets to visualize the track of electrodes (white arrow).

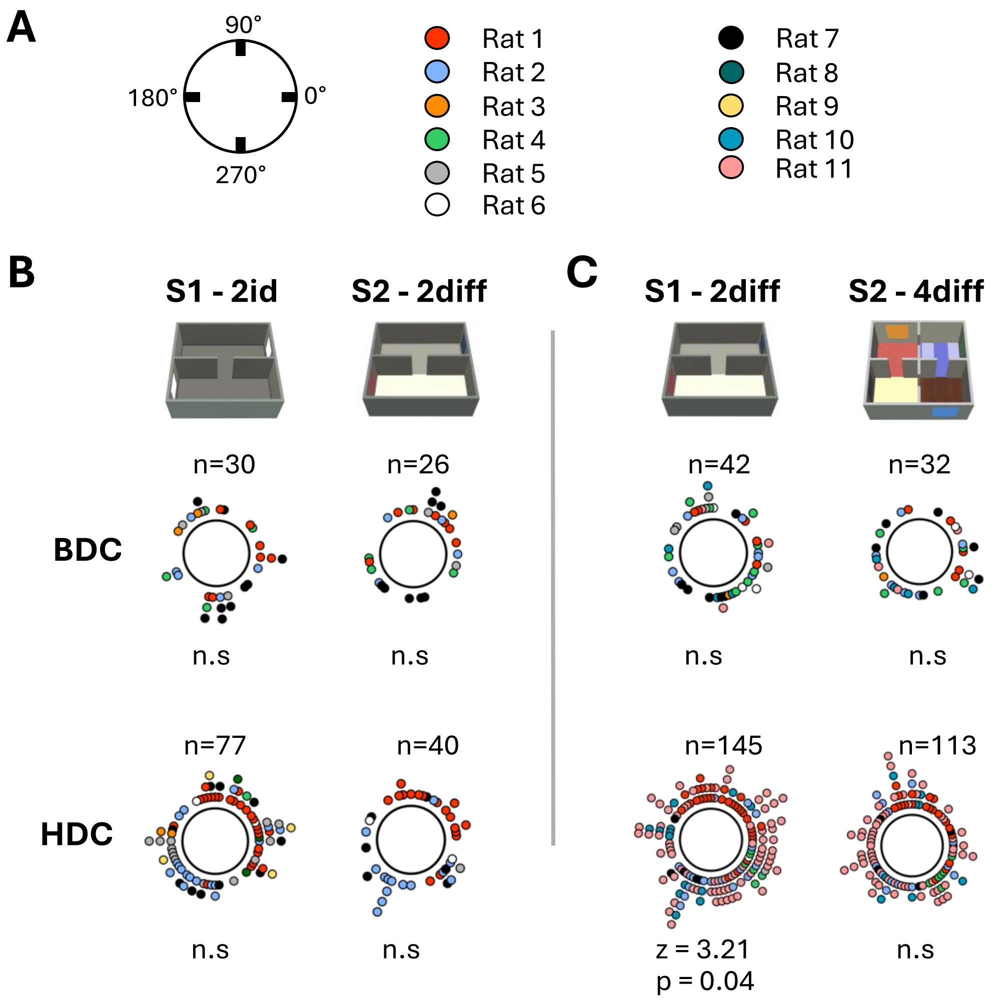


**Supplementary figure 3 – Directional firing of RSC BDC and HDC**

**A.** Left: Diagram of firing direction, Right: color code for each rat. **B.** Polar distribution of PFD for both BDCs (middle) and HDCs (bottom) was homogeneous (determined by Rayleigh test) for the 2 connected room environments. **C.** Polar distribution of PFD for BDCs (middle) was homogeneous (determined by Rayleigh test) for both the 2 and 4 connected room environments (middle) and for HDCs for the 4 connected rooms environment (bottom right), while the distribution of PFD was clustered for HDCs in the 2 different connected rooms environment (bottom left).

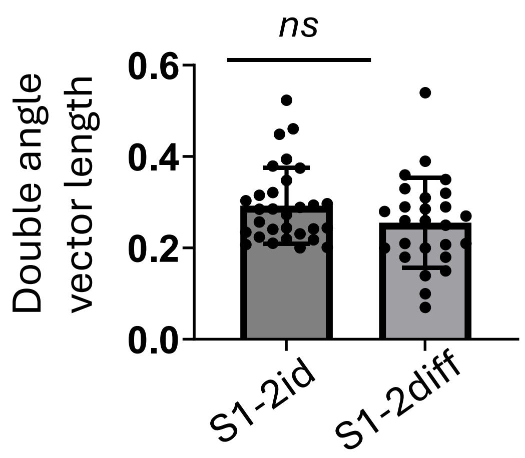


**Supplementary figure 4 – Comparison of directional activity for BDCs in 2 connected rooms environments.**

Means of Rayleigh vector length after using the double-angle procedure for BDCs directional firing (see Meethods) show no difference between 2 identical and 2 difference connected rooms environments. Bars represent mean ± SD. Statistical comparison was performed using an unpaired t-test.

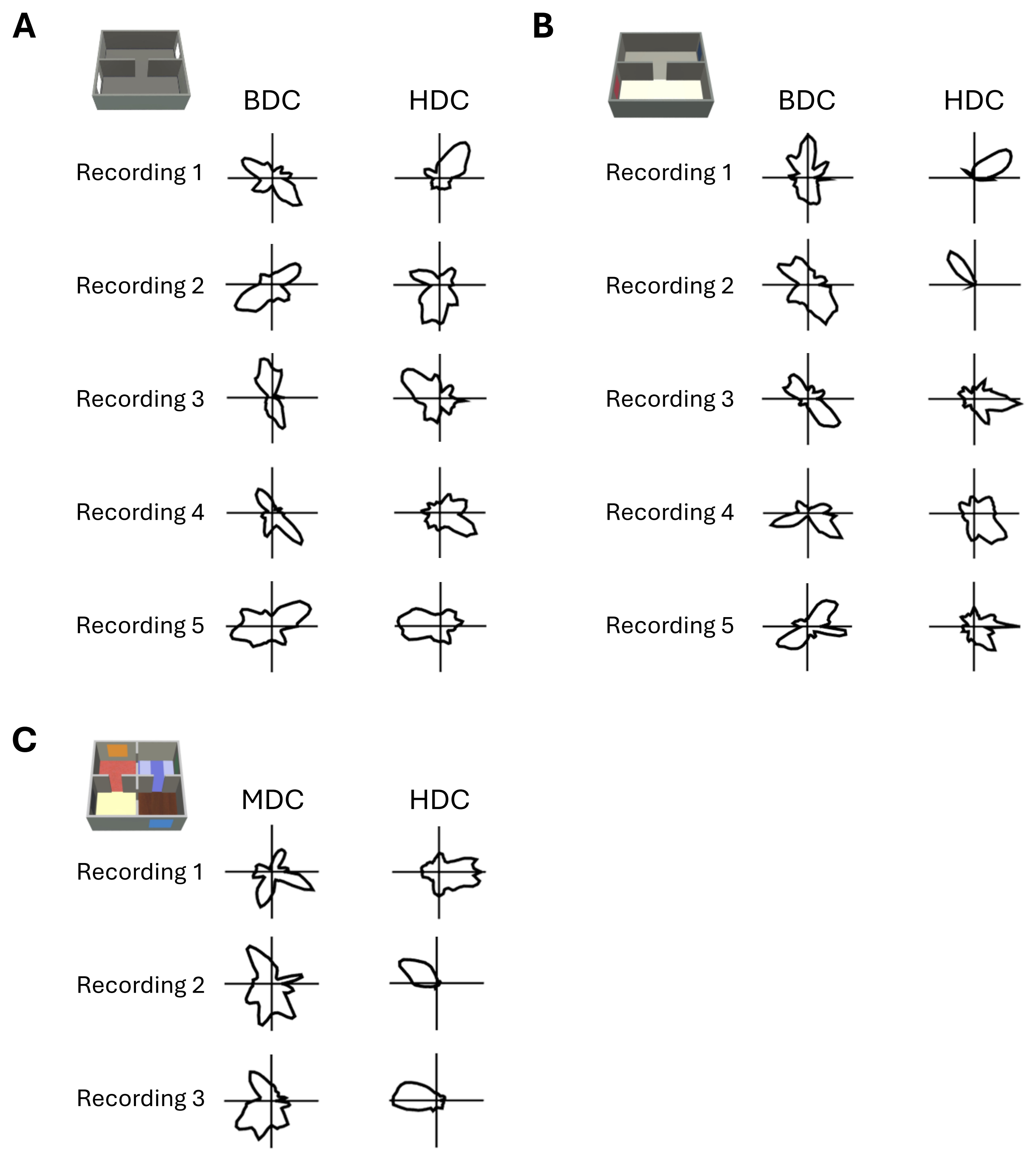


**Supplementary figure 5 – Simultaneous recording of BDCs / MDCs and HDCs.**
**A.** 5 examples from 5 different days and 4 different animals of simultaneous recordings of BDCs and HDCs in the 2 identical connected rooms environment. **B.** 5 examples from 5 different days and 3 different animals of simultaneous recordings of BDCs and HDCs in the 2 different connected rooms environment. **C.** 3 examples from 2 different days and 2 different animals of simultaneous recordings of MDCs and HDCs in the 4 different connected rooms environment.

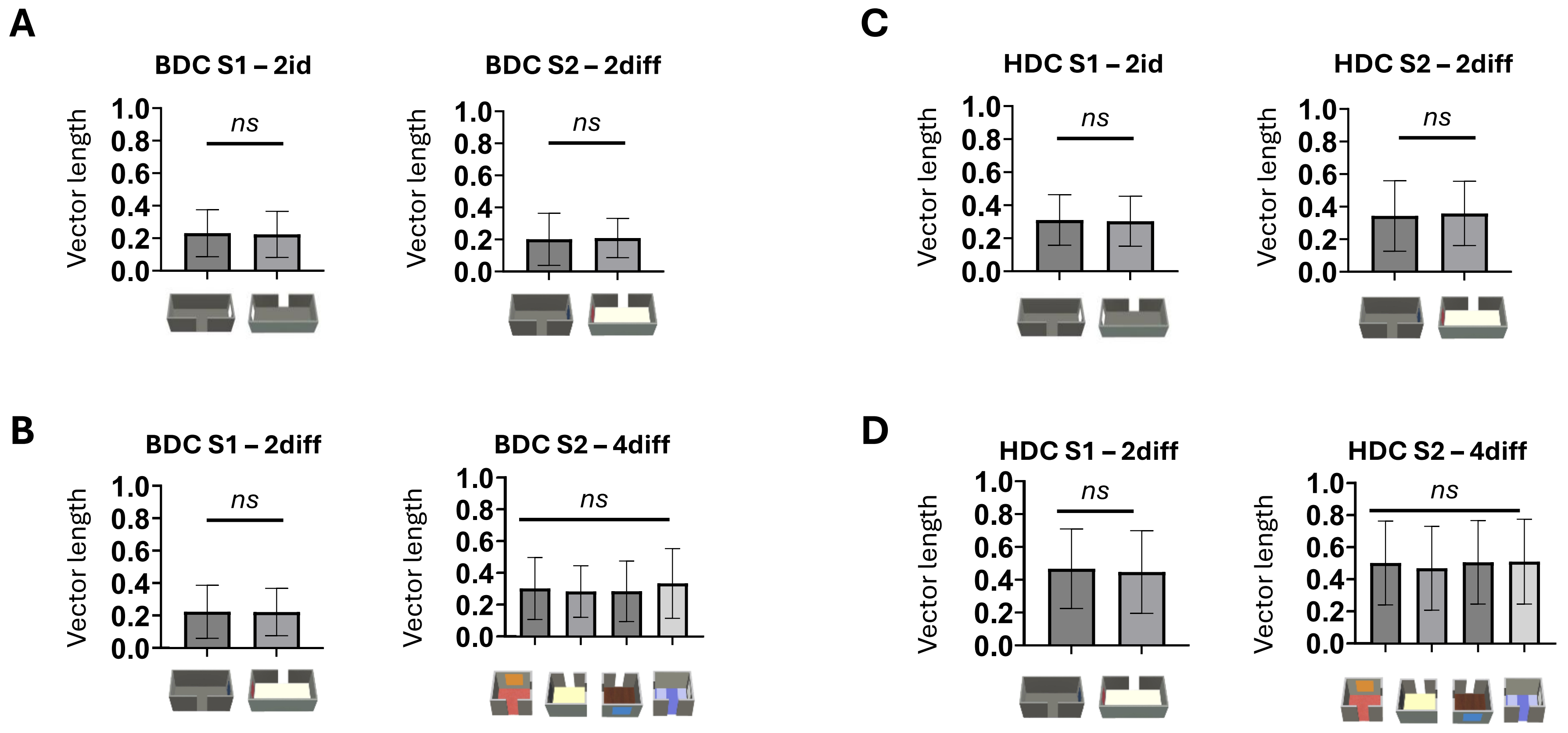


**Supplementary figure 6 – Comparison of Rayleigh vector length between rooms.**
Means of Rayleigh vector length for each room in isolation show no difference for both BDCs (**A-B**) and HDCs (**C-D**). Bars represent mean ± SD. Statistical comparison was performed using a paired t-test for 2-rooms comparisons and a one-way ANOVA for 4-rooms comparisons.

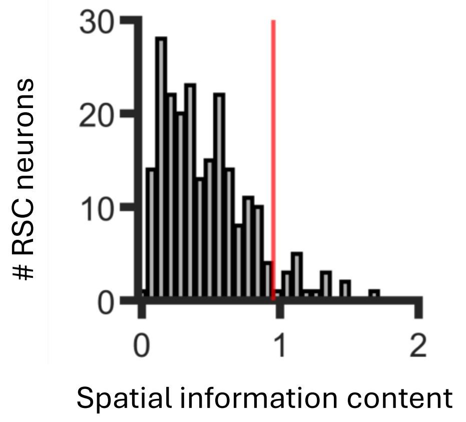


**Supplementary figure 7 – Distribution of the spatial information content of RSC cells in the 2-identical rooms environment.**
The distribution of spatial information content among non-directional RSC cells reveals a distinct threshold at 0.9, indicating a subset of neurons with significantly elevated spatial encoding.

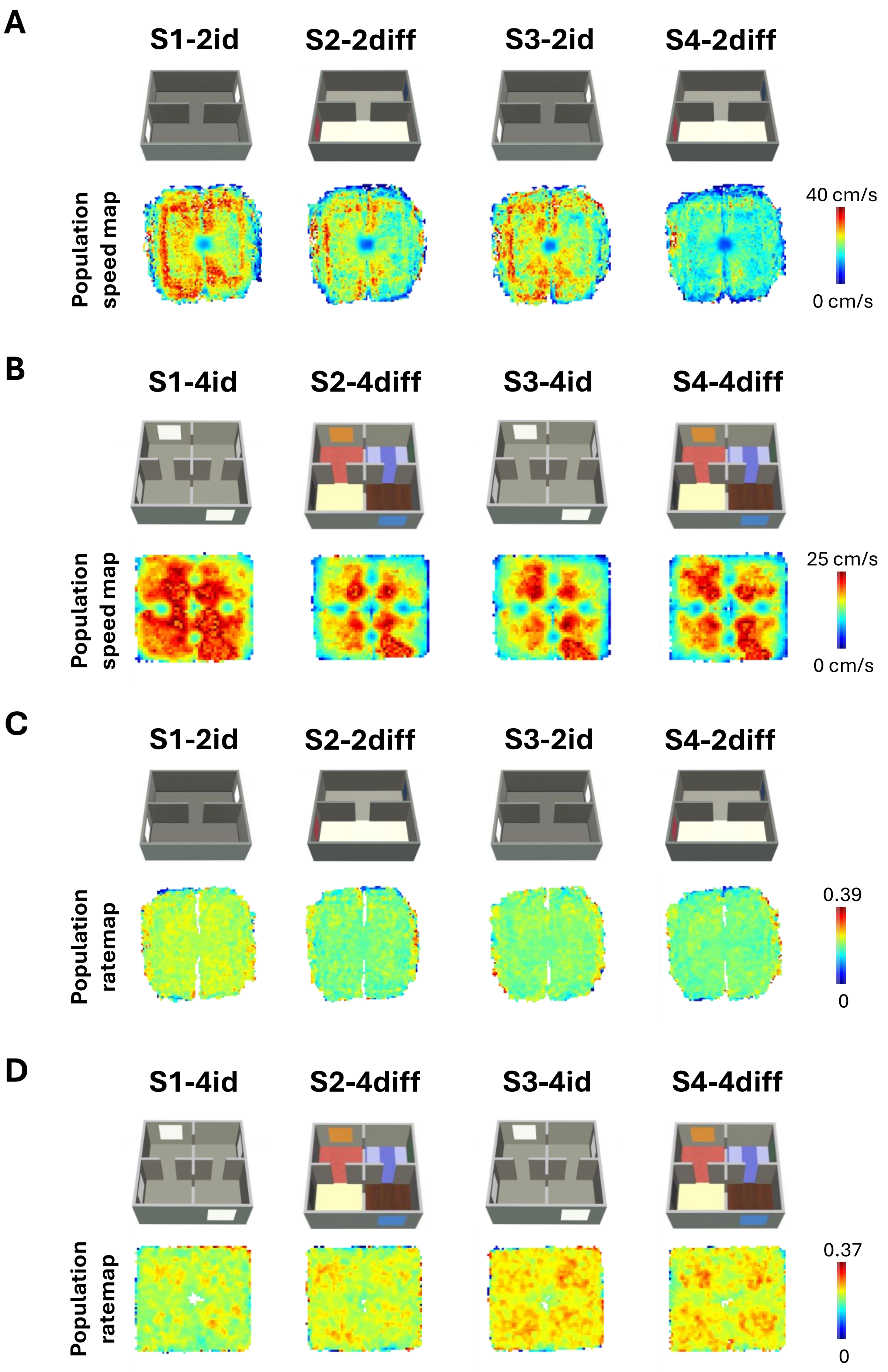


**Supplementary figure 8 – Population speed and rate maps of RSC neurons across consecutive sessions.**
**A-B.** Population maps of the average animal running speed over 35 sessions in two-room environments (A) and over 19 sessions in four-room environment (B). For each population speed map, we compare the speed observed at door(s) with the 5$^{th}$ percentile of a shuffle distribution (speed of thousand zones equal to the size of the door and located randomly in the map). For all comparison, the speed at the door fall below the 5$^{th}$ percentile of the associated shuffling analysis (speed at doors for two-room environment, S1-2id = 13.65 cm/s; S2-2diff = 9.12 cm/s; S3-2id = 10.51 cm/s; S4-2diff = 9.09 cm/s; 5$^{th}$ percentile of the shuffle distribution : S1-2id = 22.35 cm/s; S2-2diff = 17.05 cm/s; S3-2id = 18.37 cm/s; S4-2diff = 15.53 cm/s; speed at doors for four-room environments, S1-4id: top=10.63 cm/s; left=11.22 cm/s; bottom=10.66 cm/s; right=12.13 cm/s; S2-4diff: top=9.05 cm/s; left=8.70 cm/s; bottom=8.78 cm/s; right=9.20 cm/s; S3-4id: top=8.95 cm/s; left=8.64 cm/s; bottom=8.02 cm/s; right=8.04 cm/s; S4-4diff: top=8.07 cm/s; left=7.74 cm/s; bottom=7.70 cm/s; right=8.07 cm/s; 5$^{th}$ percentile of the shuffle distribution : S1-4id = 12.77 cm/s; S2-4diff = 9.65 cm/s; S3-4id = 9.33 cm/s; S4-4diff = 9.02 cm/s. Color bars indicate the highest and lowest speed. **C-D.** Population maps of the average normalized ratemaps of 222 RSC neurons recorded in two-room environments (C) and 76 RSC neurons recorded in four-room environment (B). A shuffle distribution analysis similar to population speed maps was performed to test significant changes of firing rate at doors. No observed firing rates at doors were significantly lower or greater than the 5$^{th}$ nor the 95$^{th}$ percentile of the shuffle distribution, suggesting that the spatial activity was homogeneous across the entire environments. Color bars indicates normalized firing across all neurons, from 0 to 1.

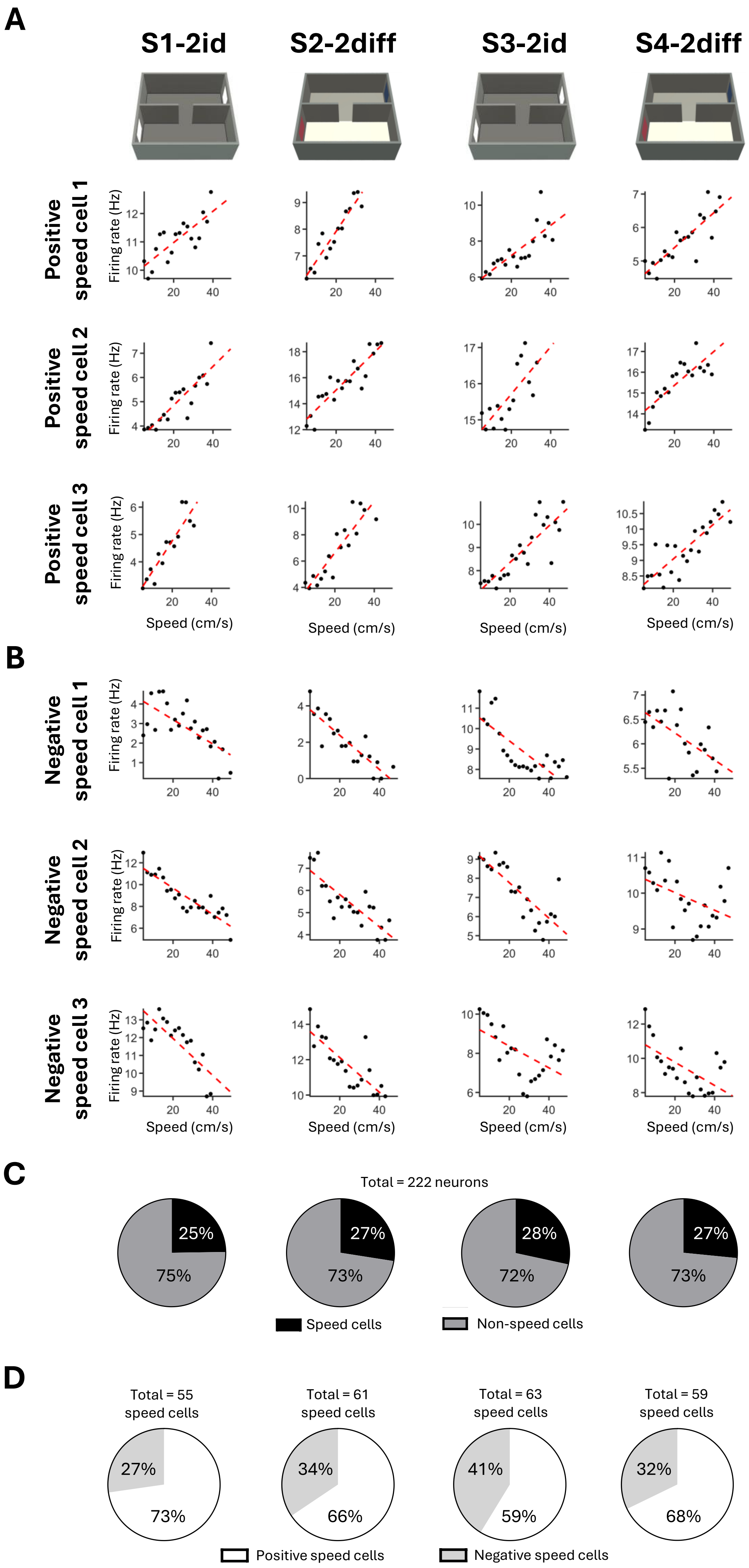


**Supplementary figure 9 – Cells with speed-modulated firing across the 2-rooms sessions.**

**A-B.** Examples of 3 RSC cells with significant positive speed-firing Pearson correlation (A) and 3 RSC cells with significant negative speed-firing Pearson correlation (B). **C.** The percentage of speed cells (black) and non–speed cells (dark gray) (in a total of 222 RSC cells) is similar between the 4 recording sessions. In speed-modulated cells, the proportion of positive and negative speed-firing modulation is also maintained between sessions.

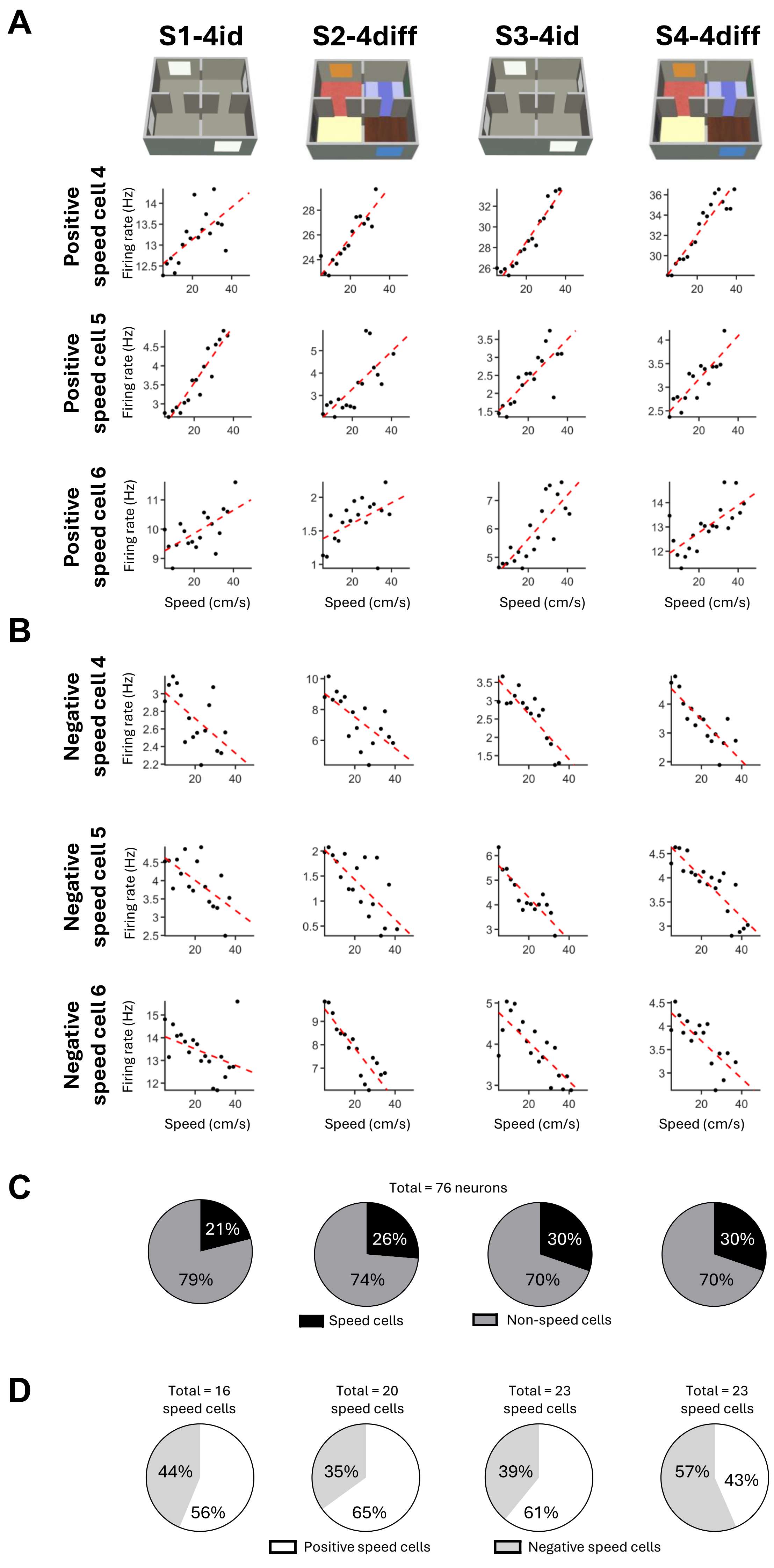


**Supplementary figure 10– Cells with speed-modulated firing across the 4-rooms sessions.**

**A-B.** Examples of 3 RSC cells with significant positive speed-firing Pearson correlation (A) and 3 RSC cells with significant negative speed-firing Pearson correlation (B). **C.** The percentage of speed cells (black) and non–speed cells (dark gray) (in a total of 76 RSC cells) is similar between the 4 recording sessions. In speed-modulated cells, the proportion of positive and negative speed-firing modulation is maintained between the 3 first sessions and change for the last session (S4-4diff).

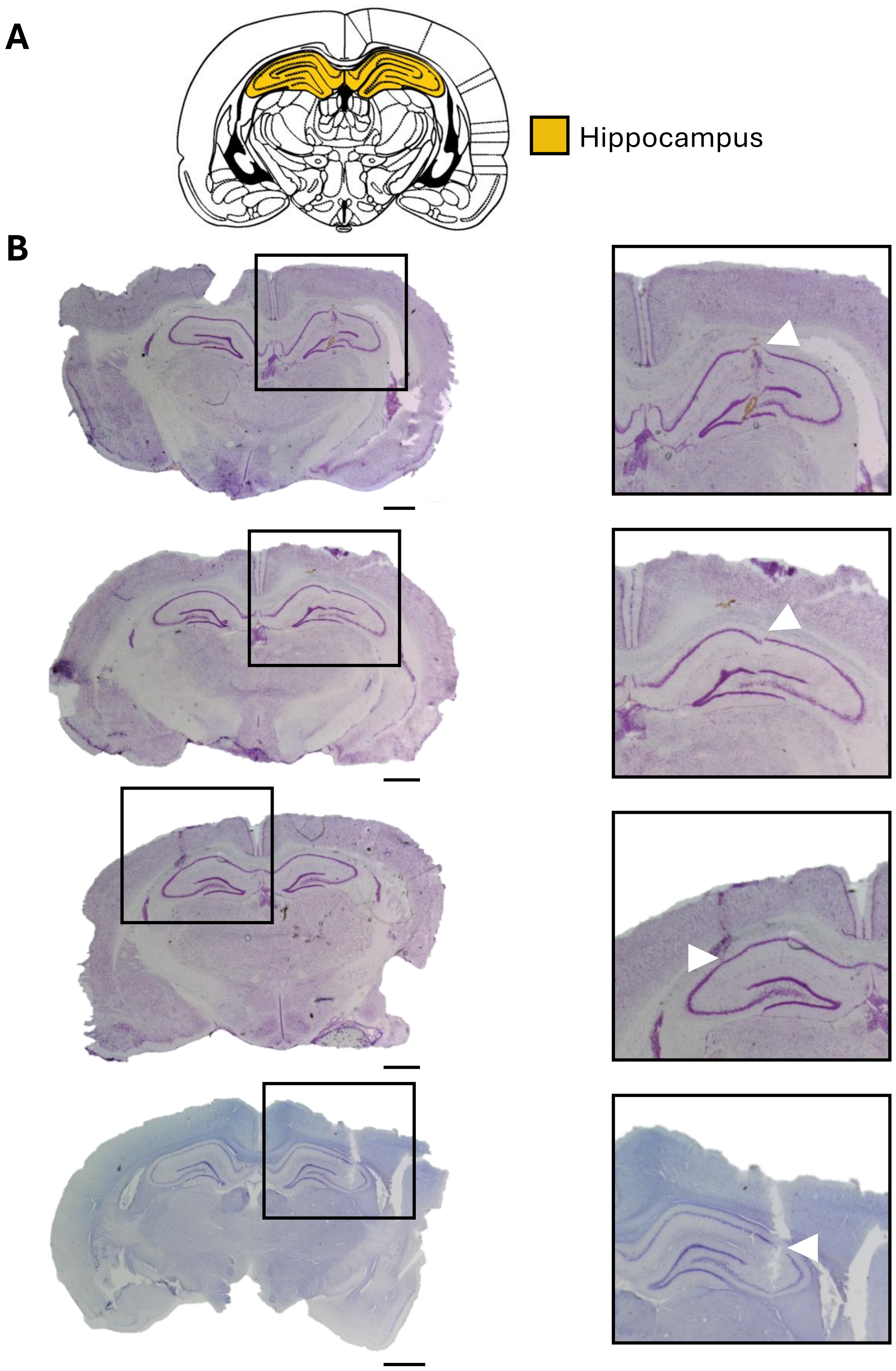


**Supplementary figure 11 – Histology hippocampus.**
**A.** Diagram showing the location of the hippocampus (HPC) adapted from Paxinos G, Watson C. The Rat Brain in Stereotaxic Coordinates. 4th Edition. Academic Press 1998. **B.** Coronal slices through the region of electrode implantation from the HPC (black insets) with scale bars indicating 1 mm. Bottom right: Magnification of insets to visualize the track of electrodes (white arrow).

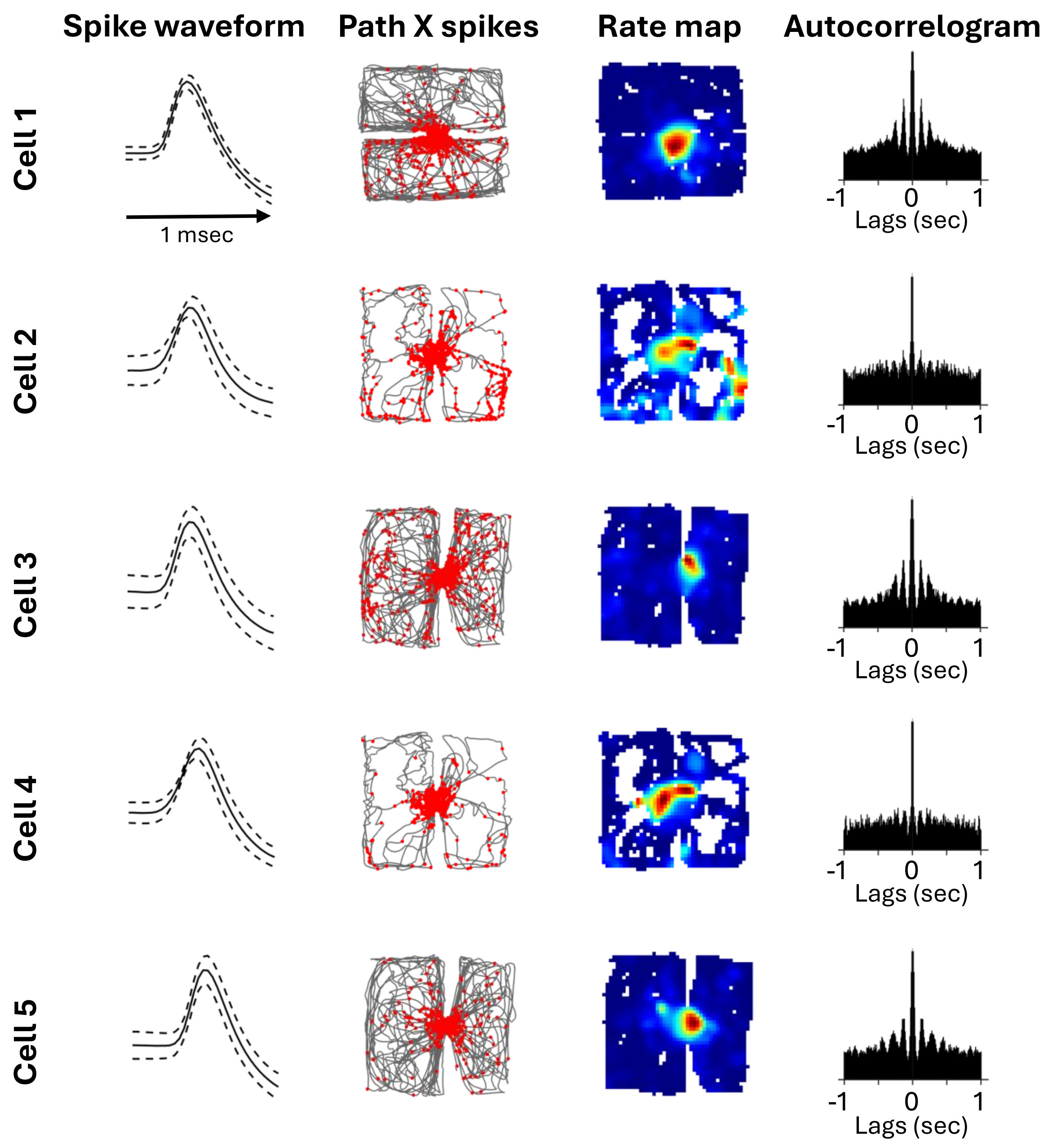


**Supplementary figure 12 – 5 place cells with a place field located at the door in the 2 connected room environment.**

The 5 place cells excluded from analyses because they have a place field located at the door in the 2 connected room environment. For each place cell is shown cell waveform (left column), animal path (grey line) with spike superimposed (red dots) (middle-left), rate maps coded according to a color scale that goes from blue (0Hz) to red (maximum rate), with cyan, yellow and orange as intermediate firing from low to high (middle-right), and autocorrelograms of the firing activity in 1-s lags (right).

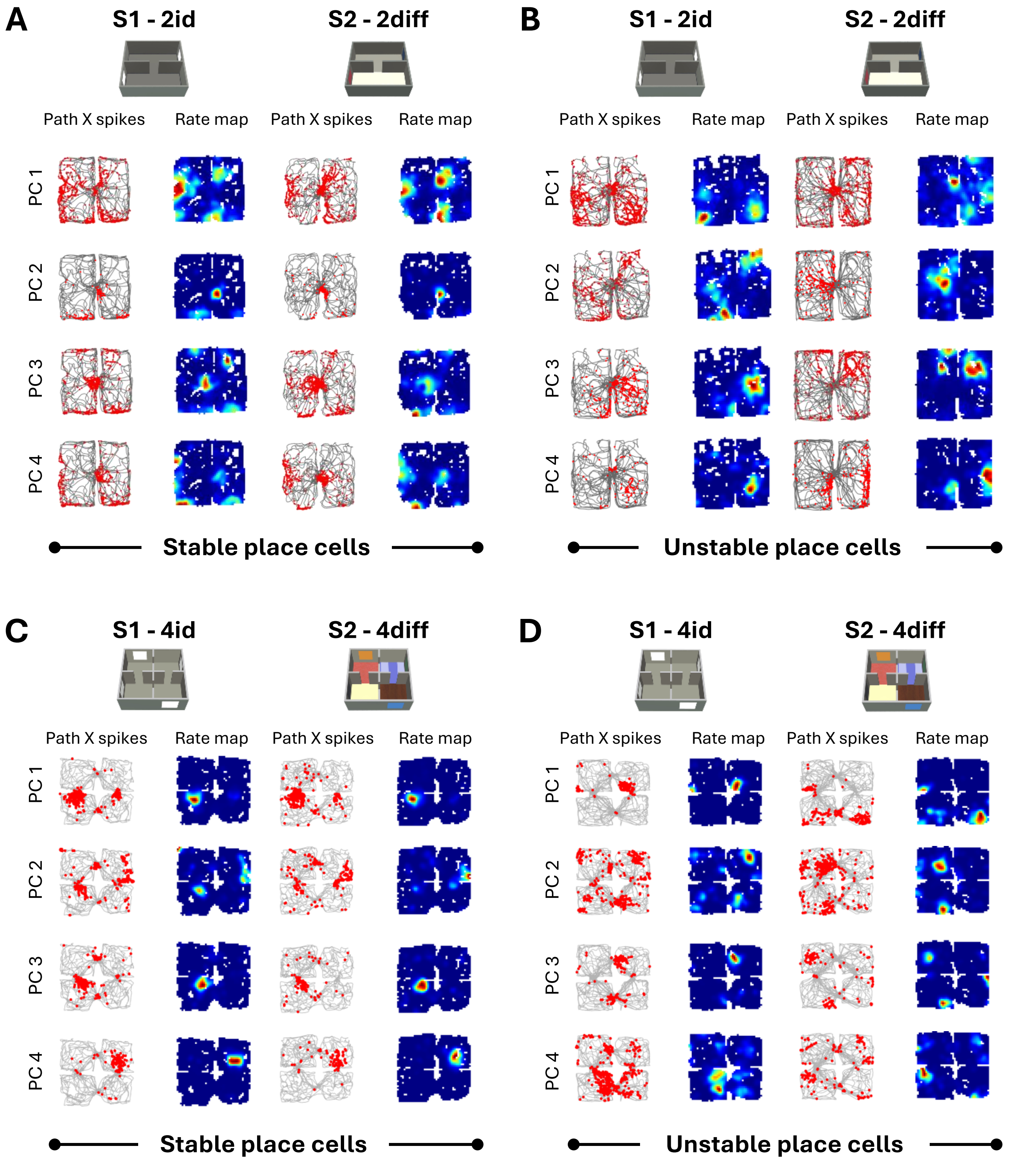


**Supplementary figure 13 – Place cells (PC) stability between identical and different connected rooms environment is session dependent.**
Examples of the activity of simultaneously recorded PC between identical (left) and different (right) connected rooms environment. Within a given session PCs show either stable place field(s) (**A-C**) and unstable place field(s) (**B-D**). For each place cell is shown animal path (grey line) with spike superimposed (red dots) (left) and rate maps coded according to a color scale that goes from blue (0Hz) to red (maximum rate).
